# Structure of a rabies virus polymerase complex from electron cryo-microscopy

**DOI:** 10.1101/794073

**Authors:** Joshua A. Horwitz, Simon Jenni, Stephen C. Harrison, Sean P.J. Whelan

**Author notes:** Correspondence: Stephen C. Harrison, Sean P.J. Whelan. Lead contact: Stephen C. Harrison.

## Abstract

Non-segmented negative-stranded (NNS) RNA viruses, among them the virus that causes rabies (RABV), include many deadly human pathogens. The large polymerase (L) proteins of NNS-RNA viruses carry all the enzymatic functions required for viral mRNA transcription and replication: RNA polymerization, mRNA capping, cap methylation. We describe here a complete structure of RABV L bound with its phosphoprotein cofactor (P), determined by electron cryo-microscopy at 3.3 Å resolution. The complex closely resembles vesicular stomatitis virus (VSV) L-P, the one other known full-length NNS-RNA L protein structure, with key local differences (e.g., in L-P interactions). Like the VSV L-P structure, the RABV complex analyzed here represents a pre-initiation conformation. Comparison with the likely elongation state, seen in two partial structures of pneumovirus L-P complexes, suggests differences between priming/initiation and elongation complexes. Analysis of internal cavities within RABV L suggests distinct template and product entry and exit pathways during transcription and replication.

## INTRODUCTION

Rabies virus (RABV) and other viruses with non-segmented, negative-strand (NNS) RNA genomes have, as the catalytic core of their replication machinery, a large, multifunctional RNA polymerase (L). Many of these viruses are serious human pathogens, including Ebola virus, respiratory syncytial virus, and measles. Vaccines are available for some, including descendants of the storied work on rabies by Pasteur and Roux, but specific, small-molecule therapeutics are still under development. By analogy with inhibitors of viral polymerases of many other types, L would be a suitable candidate for inhibitor development, for which structural and mechanistic studies are essential precursors.

For all mononegaviruses, including the rhabdoviruses RABV and vesicular stomatitis virus (VSV), L associates with a cofactor known as P (for phosphoprotein). P bridges L with the viral nucleocapsid, an antisense RNA genome fully encapsidated by the viral nucleoprotein (N). The full-length structure of L from VSV (Liang et al., 2015), the sole example published so far of a complete NNS viral polymerase, shows the global features that sequence comparisons suggest it has in common with most other NNS RNA viral L proteins. Its multidomain organization includes three enzymatic modules: an RNA-dependent RNA polymerase (RdRp), a capping domain (CAP), and a dual-specificity methyltransferase (MT) domain. CAP is a GDP:polyribonucleotidyltransferase (PRNTase) that transfers a 5’ monophosphate of the nascent RNA transcript onto a GDP acceptor (Ogino and Banerjee, 2008); the single MT domain methylates the ribose 2’-O position on the first nucleotide of the transcript and then the N-7 position of the capping guanylate (Rahmeh et al., 2009). Transcription initiation and cap addition require several residues within or proximal to a priming loop from CAP, which extends into the RdRp core. Residues at related positions in the RABV L amino-acid sequence are similarly essential for its activities. A connector domain (CD) and a C-terminal domain (CTD), both with non-enzymatic functions, flank the MT domain and, in conjunction with P, likely facilitate the large structural rearrangements that appear to coordinate the three enzymatic activities during transcription and replication.

We report here an atomic model of L from RABV SAD-B19 in complex with a 91-residue, N-terminal fragment of P (P_1-91_), from electron cryo-microscopy (cryo-EM) at an average resolution of 3.3 Å. The atomic structure resembles that of VSV L-P in many respects, consistent with the 34.1% amino acid sequence conservation between the two L proteins. As in the VSV L-P complex, binding of P_1-91_ locks the CD, MT domain, and CTD into a fixed, “closed” arrangement with respect to the large RdRp-CAP module. This closed conformation appears to represent the L protein poised for initiation at the 3’ end of the genome or anti-genome. Comparison with structures of two recently published pneumovirus L-P complexes (Gilman et al., 2019) (Pan et al., submitted) as well as with that of a VSV L-P reconstruction determined at 3.0 Å resolution (Jenni, et al., submitted) suggests that replication and transcription have alternative priming configurations and alternative product exit sites.

## RESULTS AND DISCUSSION

### Identification of a stable L-P complex

We have determined previously that a minimal fragment of the N-terminal domain of RABV P, P_11-50_, is sufficient to stimulate processivity of RABV L on a non-encapsidated RNA template derived from the 3’ leader sequence of the RABV genome (Morin et al., 2017). In our prior studies of VSV, we found a minimal fragment of VSV P that stimulated VSV L activity in vitro and also stably anchored its C-terminal globular domains (CD, MT and CTD) against the N-terminal RdRP and CAP domains (Rahmeh et al., 2012; Rahmeh et al., 2010). We visualized RABV L alone or in complex with various fragments of P by negative-stain electron microscopy (Fig. 1) and found that in the absence of P, the C-terminal domains of RABV L, like those of VSV L, have a range of positions with respect to the ring-like RdRp-CAP core (Fig. 1B). While P_11-50_ stabilized the C-terminal domains of RABV L, a longer fragment (P_1-91_) shifted their positions further, to a degree equivalent to that achieved by adding full-length P (P_FL_) (Fig. 1C). To fit the structure of VSV L (obtained in complex with VSV P_35-106_) into a low-resolution map for RABV L-P_11-50_, we needed to fit the C-terminal domains at an ∼80° offset (data not shown), but we could readily fit the low-resolution maps of RABV L-P_1-91_ and RABV L-P_FL_ with the structure of VSV L as a single rigid body (Fig. 1C, right). We inferred that the RABV L-P_1-91_ and VSV L-P_35-106_ complexes reflected analogous states and therefore used P_1-91_ to determine the RABV L structure by cryo-EM.

**Figure 1:**
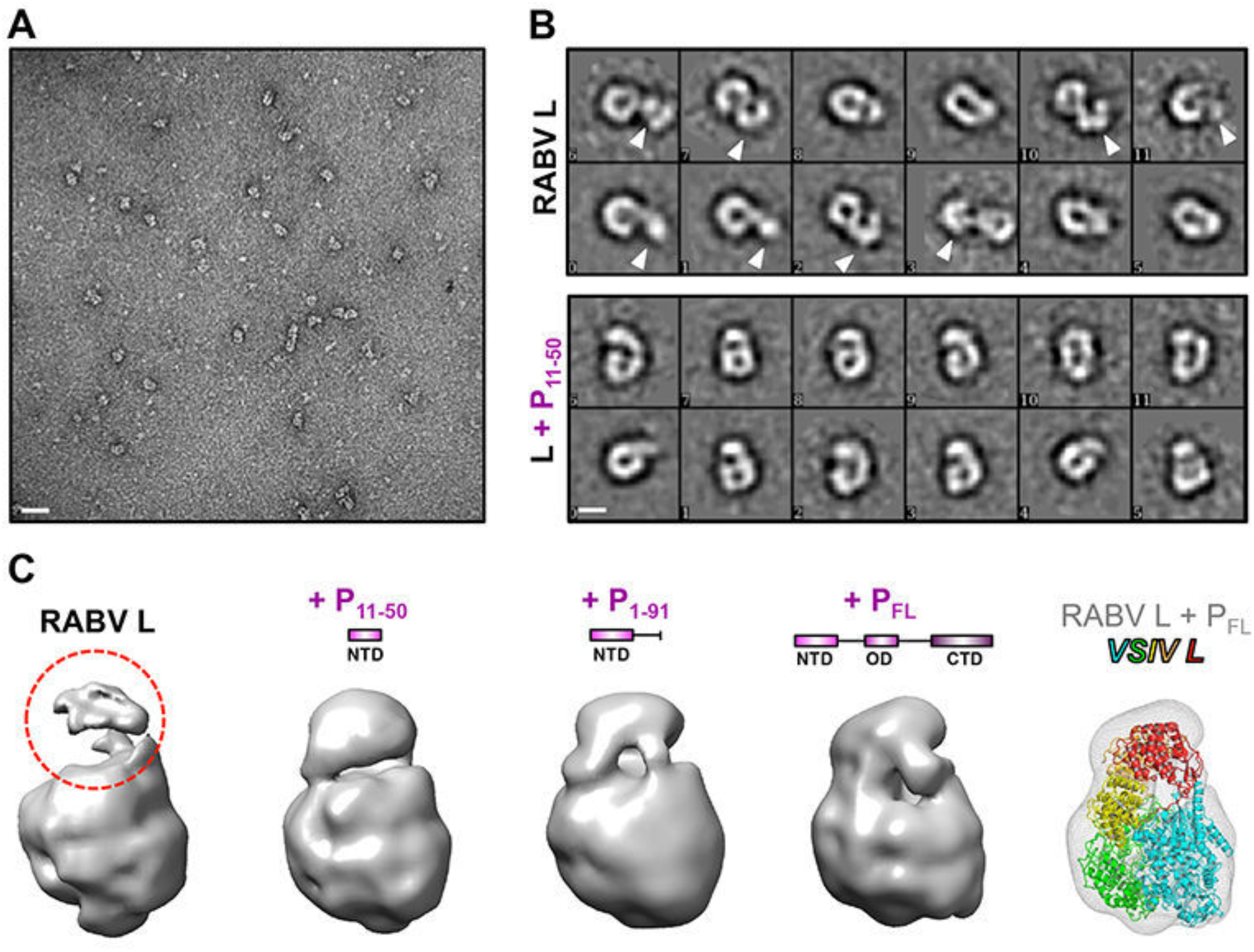
Effect of P on conformation of L. (A) Electron micrograph of negatively stained RABV L. Scale bar: 25 nm. (B) Representative 2D class averages of L alone or in complex with P_11-50_. Scale bar: 5 nm. Arrows indicate flexible conformations of the three C-terminal globular domains of L. (C) 3D reconstructions of negatively stained L alone or in complex with the indicated fragment of P. Red dashed circle indicates poor density for the disordered C-terminal globular domains in the absence of any P. *Right,* the VSV L atomic model fit into the RABV L-P_FL_ reconstruction.

### Cryo-EM structure determination

We immobilized purified complexes of RABV L-P_1-91_ in vitreous ice and visualized them by cryo-EM (Fig. S1A, B) as described in Methods. We created an initial 3D reference *de novo* using the ab-initio module in cisTEM (Grant et al., 2018) from dose-fractionated movies obtained at 200 kV acceleration, and obtained a 6.7 Å resolution map from 22,000 particles after classification and refinement using RELION (Scheres, 2012) (Fig. S1C). In extending the resolution from movies recorded at 300 kV acceleration, preferred orientation of the particles in the ice complicated particle selection and 3D reconstruction. By using an angular binning procedure to decrease representation of common views (Fig. S1D and S2), we obtained a reasonably isotropic reconstruction using cisTEM. After per-particle, projection-based CTF and motion correction (Bayesian polishing) in RELION 3.0 (Nakane et al., 2018; Zivanov et al., 2018), refinement in cisTEM yielded a map with an average resolution of 3.3 Å (FSC=0.143 criterion; Figs. 2A, S1E and S2).

**Figure 2:**
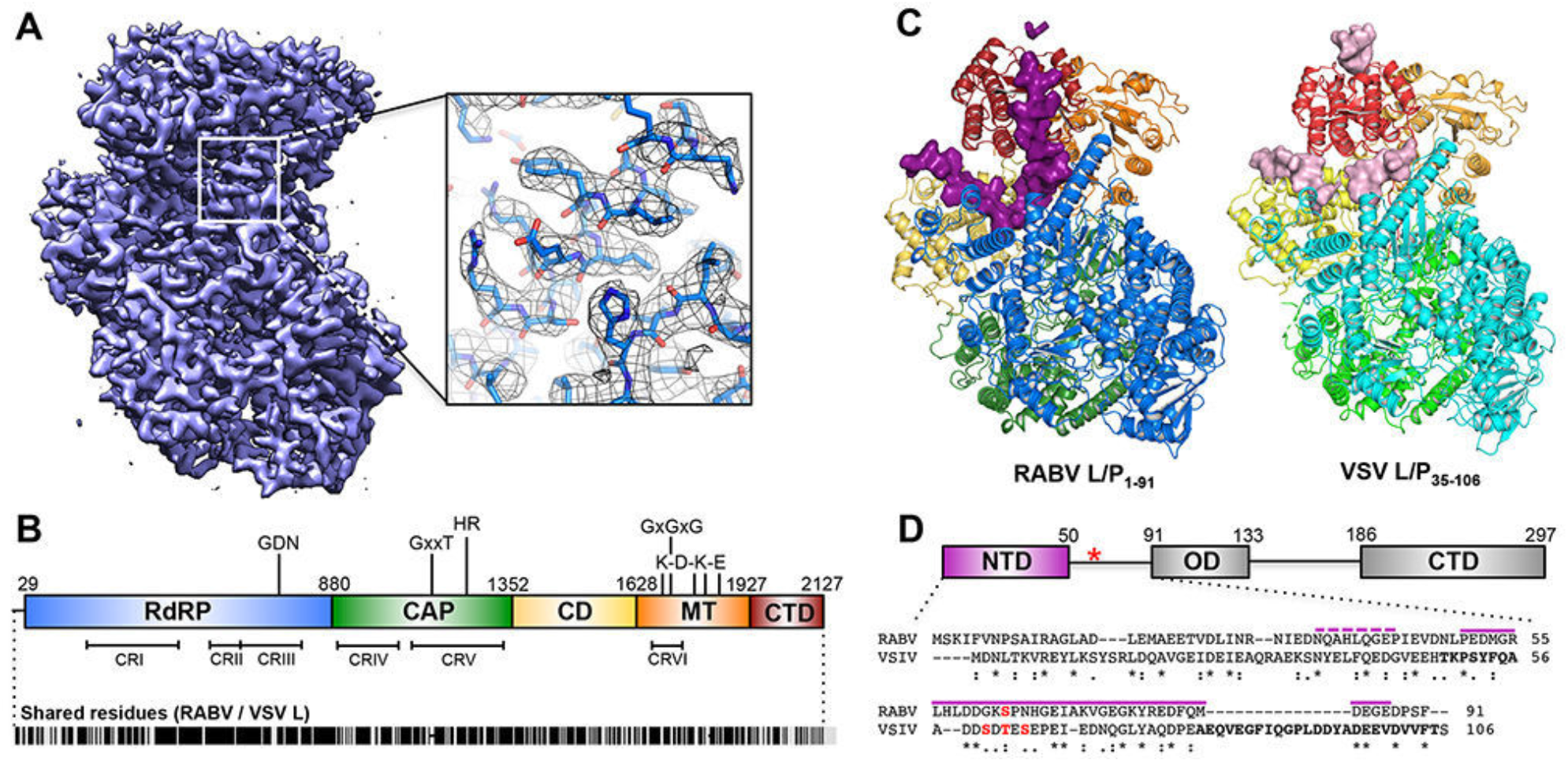
Structure of the RABV L-P complex. (A) 3.3 Å cryo-EM map of RABV L-P_1-91_ and model fit into the density, with clear side-chains even on exterior residues. (B) Domain organization of RABV L, with key catalytic residues listed. Numbers give amino acid positions at domain boundaries. Below, locations of conserved regions (CRI-VI) throughout NNS-RNA virus L proteins and amino-acid similarity score with VSV L (Indiana strain); black lines indicate identical residues (∼35%). (C) Structure of RABV L-P_1-91_ (left) and the model of VSV L-P_35-106_ from a 3.0 Å cryo-EM map (right) (Jenni, et al.). Domains are colored as in (B). P for both structures is shown in surface representation, colored in plum (RABV) and light pink (VSV). The five unassigned residues of RABV P are represented in plum ribbon. (D) Domain organization of RABV P (top, numbers as in [B]; red asterisk indicates the position of Ser63P, typically phosphorylated). Below, alignment with VSV P (purple lines, modeled RABV P residues; bold letters, modeled VSV P residues; asterisks, shared residues; colons, chemically similar residues; periods, weakly similar residues; red, phosphorylation sites).

To generate an atomic model, we aligned and threaded the sequence of RABV L onto the atomic coordinates of VSV L and crudely fit the model to the map using Rosetta (Rohl et al., 2004a; Rohl et al., 2004b; Song et al., 2013). After manual adjustment, we could fit nearly all of the 2127-residue polypeptide chain of L from residue 29 to the C-terminus (Fig. 2). We assigned residues for regions of continuous, but disordered, density in the CAP priming loop (1177-84) and the long linker between the CD and MT domains (1624-35), because there was little ambiguity about the general course of the polypeptide chain, but the Cα positions and the side chain orientations are imprecise. Several inter-domain linkers that were ambiguous in the VSV L map have continuous and ordered density for the analogous regions in the RABV L map, permitting assignment of their residues with high confidence. The clarity of side-chains throughout all five domains of L indicates that regions of the map with poor density reflect local flexibility rather than wholesale domain movements (Fig. S3). Domain-specific refinement from a much larger portion of the dataset using RELION Multi-Body showed significant inter-domain flexibility (Fig. S4) and suggested that the comparatively small fraction of the dataset used for the 3.3 Å reconstruction reflects a particular configuration within a somewhat broader ensemble.

Rigorous selection of this small set of homogeneous particles for the 3.3 Å map yielded clear and nearly continuous density for the C-terminal half of P_1-91_. Visible density for some side-chains on a stretch of P bound to the CTD permitted assignment of the sequence register in this region, from which we have inferred the approximate residue positions for the remainder of visible P density. We have therefore assigned 37 of the 91 residues of P included in the complex (Fig. 2C,D), not including a stretch of 5 residues, which we could not assign, bound to the top of the CTD. The cryo-EM data collection and model validation statistics for the map and complete RABV L-P structure are summarized in Table S1.

**Table 1.** Cryo-EM data collection and validation statistics.

| ***Data collection and processing*** |  |
| --- | --- |
| Microscope | Tecnai F30 Polara |
| Detector | Gatan K2 Summit |
| Magnification | 31,000X |
| Pixel size (Å) | 1.234 |
| Voltage (kV) | 300 |
| Exposure (e<sup>-</sup>/Å<sup>2</sup>) | 72 |
| Defocus range (µm) | 0.6-3.5 |
| Final particles | 44,500 |
| Symmetry | C1 |
| Resolution (Å) | 3.29 |
| ***Model composition, refinement and validation*** |  |
| Non-hydrogen atoms | 17219 |
| RMSD bonds (Å) | 0.005 |
| RMSD angles (°) | 1.345 |
| Ramachandran: |  |
| Favored (%) | 99.44 |
| Allowed (%) | 0.47 |
| Outliers (%) | 0.09 |
| Rotamer outliers (%) | 0.00 |
| Clash score | 6.65 |
| MolProbity score | 1.37 |
| ***Model-to-map fit*** |  |
| Cross-correlation coefficient (mask) | 0.75 |
| Cross-correlation coefficient (volume) | 0.76 |
| Main-chain | 0.76 |
| Side-chain | 0.71 |

The structures of RABV and VSV L in complex with their respective P fragments are very similar (Fig. 2C). Rigid-body alignment of their structures yields a rmsd of 1.905 Å based on 1539 Cα residues; individual domain rmsds are 0.905 Å (595 Cα), 1.071 Å (352 Cα), 2.088 Å (186 Cα), 1.504 Å (233 Cα), and 3.675 Å (120 Cα) for the RdRp, CAP, CD, MT, and CTD domains superposed independently (Fig. S5). There is nearly complete conservation of the secondary structure elements from end to end, owing to strong sequence conservation between each of the five domains for both proteins: shared amino acid identities are 36.1%, 41.9%, 25.4%, 31.2%, and 20.1% for the RdRp, CAP, CD, MT, and CTD domains, respectively. The overall course of P as it transits through L is also similar for both RABV and VSV, despite broad differences in sequence among the residues that could be assigned for the respective P fragments, and the mere 11.8% shared amino acid identities between them overall.

### Phosphoprotein fragment P_1-91_

We detect two segments of density for the P_1-91_ polypeptide chain (Fig. 3). One segment binds in a groove on the outward-facing surface of the CTD above the C-terminal residues of L (Fig. 3A, panel i). It is too short to assign its sequence with confidence, but biochemical experiments that measured RABV L processivity on naked RNA templates in the presence of N-terminal fragments of P suggest that this interaction involves at least some of the residues immediately following Gln40 of P (Morin et al., 2017). The second segment is much longer, with a continuous stretch of 37 residues. It contains about 11 residues bound on the proximal face of the CTD, probably in part through a salt bridge between Glu51P and Arg2039L (Fig. 3A, panel ii), about 3 residues in contact with a long RdRp helix (residues 746-773), probably including a salt bridge between Asp60P and Arg754L (Fig. 3A, panel iii), and about 13 additional residues in the groove between the RdRp and CD (Fig. 3A, panel iv). The C-terminus of the ordered segment we could build is Glu87P. The closer association of P with the proximal face of the CTD of L for RABV relative to VSV suggests RABV P partially compensates for the C-terminal tail that is “missing” in RABV L (Fig. 3B).

**Figure 3:**
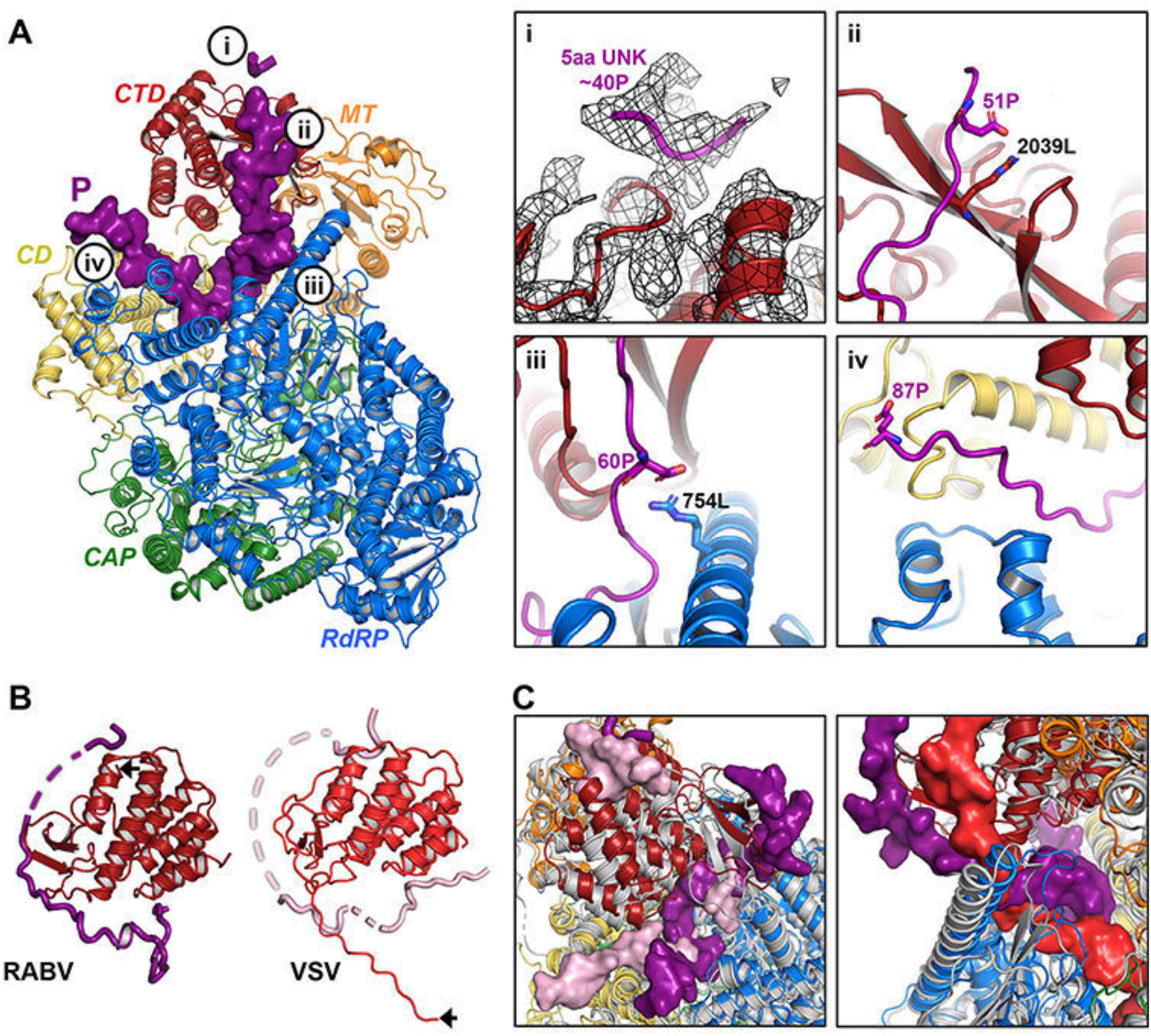
Association of RABV P_1-91_ with L. (A) *Left,* RABV L-P_1-91_ as in Fig. 2C. Bubbled letters indicate the regions shown in greater detail in panels i-iv on the right: *Right*: (i) Unassigned residues of P at the top of the CTD, with the 3.3 Å map in black mesh; (ii-iv) Indicated residues of L (black) and P (purple) in stick representation. (B) CTDs of RABV and VSV L; P is in plum for RABV and light pink for VSV. Black arrows indicate the C-terminal residues of each L. (C) Superposition of RABV L (colored cartoon, as in Fig. 2) and VSV L (gray cartoon). *Left:* different paths taken by RABV P (plum, surface) and VSV P (light pink, surface) through L. *Right:* RABV P (plum, surface) is partially coincident with the VSV C-tail (red, surface) in the inter-domain cavity of L, while VSV P occupies far less of the same cavity.

As a first check on the polarity we assigned to P, we fused GFP to the N terminus of P_1-91_ and visualized its complex with L by negative-stain EM (Fig. 4A). A low-resolution 3D reconstruction showed diffuse density for the globular GFP at the top of L, radiating from a position near the top of the long CD:MT linker (Figs. 4A, S6 and S7). This result is consistent with our assignment of the direction of P residues in the density and with our assignment of residues near Gln40P to the segment bound on the CTD outward-facing surface. The map for the L-GFP-P_1-91_ complex otherwise resembled that of the L-P_1-91_ complex without GFP (Fig. S7), indicating that N-terminal fusion of GFP to P_1-91_ had not disrupted its binding to L.

**Figure 4:**
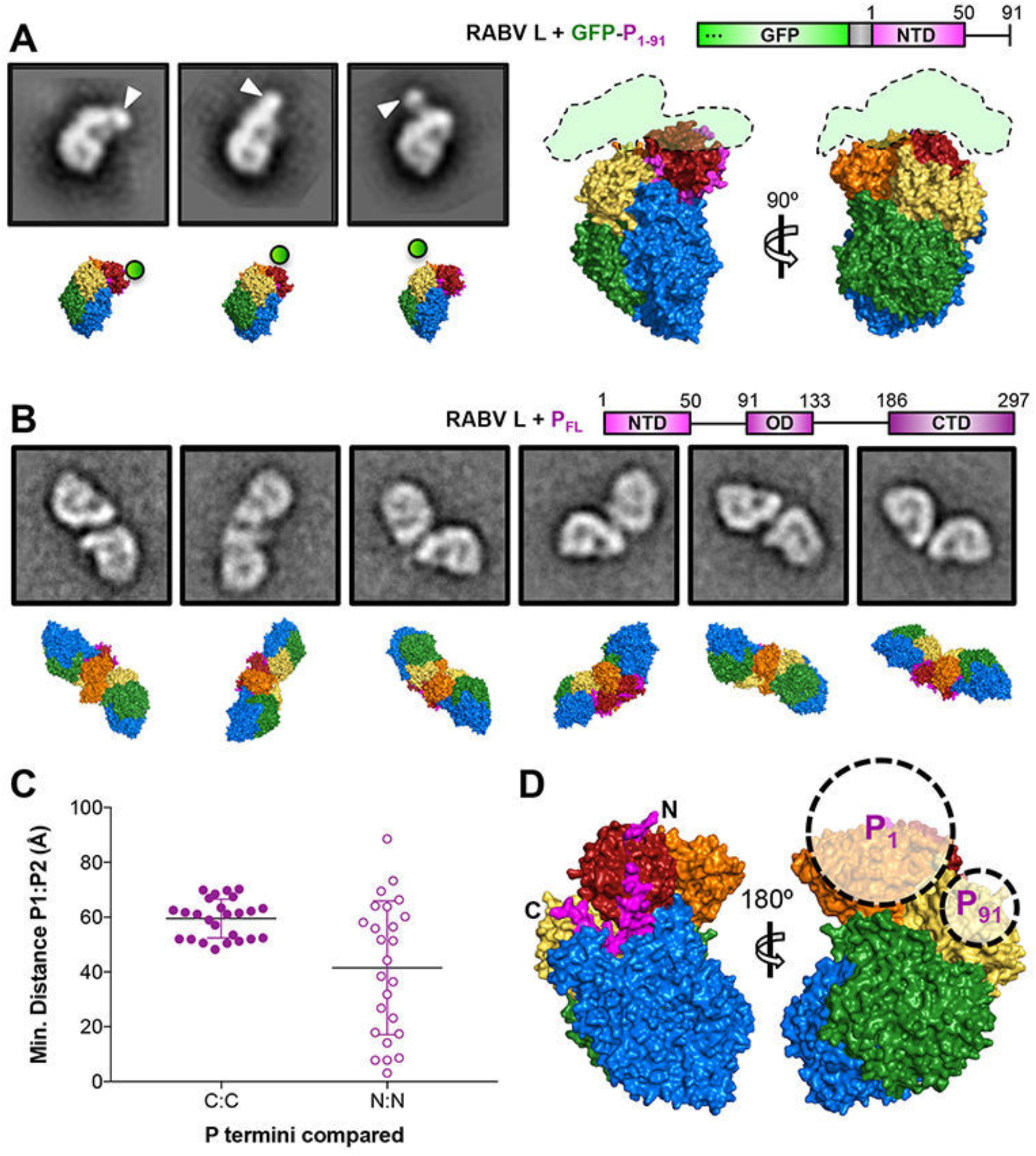
Analysis of negatively stained L-GFP-P_1-91_ and L-P dimers. (A) *Left,* 2D class averages of negatively stained L-GFP-P_1-91_ and matched projections of the RABV L atomic model with GFP position approximated (see also Fig. S6). *Right*, GFP trace from a 3D reconstruction of negatively stained RABV L-GFP-P_1-91_ (Fig. S7) superposed on the RABV L atomic model. (B) 2D classes of negatively stained RABV L-P_FL_ dimers and matched projections of the RABV L atomic model (see also Fig. S9 and S10), as described in Fig. S8. (C) Minimal distances between Cα atoms of either P_87_ (C:C) or the N terminus of the P segment bound at the CTD apex (N:N) for each projection-matched dimer. (D) Approximate locations of P_1_ and P_91_ with respect to L determined from the projection matching analysis.

We confirmed the location of the C terminus of P_1-91_ by visualizing RABV L in complex with full-length P (P_FL_) by negative-stain EM, which showed dimers of RABV L that closely resembled those of VSV L complexed with full-length VSV P (Rahmeh et al., 2012). The central, oligomerization domain of RABV P (P_OD_) includes residues 91-133, which form a homodimer (Ivanov et al., 2010). We matched projections of the high-resolution RABV L-P_1-91_ map to each L monomer in the negative-stain EM 2D class averages (Figs. 4B and S8-10). The distance between P_91_ residues in the crystal structure of the homodimeric RABV P_OD_ is 25 Å, and the last ordered residue in our model is 87, giving a spacing (assuming a flexible but extended chain for 87-91) of about 55 Å for the exit points of P on two L molecules in the L-P_FL_ dimer. Projection matching of the dimer partners in negative-stain EM images yielded spacings of ∼60Å +/-10Å, while the distances between the N termini of the short, bound segment of P varied widely from 0 to 90 Å (Fig. 4C). We also found several different dimer arrangements (Fig. S10), all compatible with the polarity of the segments of P we have modeled, with our assignment of the C-terminal residues of P_1-91_, and with flexibility of P residues 87-91.

### RNA-dependent RNA polymerase

The RdRp domains of RABV and VSV L have nearly coincident three-dimensional structures, as expected from their amino-acid sequences, which are identical at 36.1% of the aligned positions (Fig. S5). They have the familiar finger-palm-thumb structure at their core, augmented by a substantial N-terminal subdomain and by a C-terminal transition into the capping domain (Fig. S11). The catalytic site (GDN motif) is within a central hollow, connected to the exterior of the molecule by four channels. Comparison with dsRNA polymerases, for which structures of initiating and elongating complexes are known (Lu et al., 2008; Tao et al., 2002), has allowed assignment of these channels to the four molecular-passage functions required for RdRp activity: template entry, template exit, nucleotide substrate entry, and product exit (Fig. 5 and (Reguera et al., 2016)). In the initiation-competent configuration of RABV L we describe here, the product exit channel is occluded by a retractable priming loop from CAP (Fig. 5B, discussed below), while the putative template exit channel appears “closed” by a grouping of small loops that may be pushed open by the exiting template RNA (Fig. 5C).

**Figure 5:**
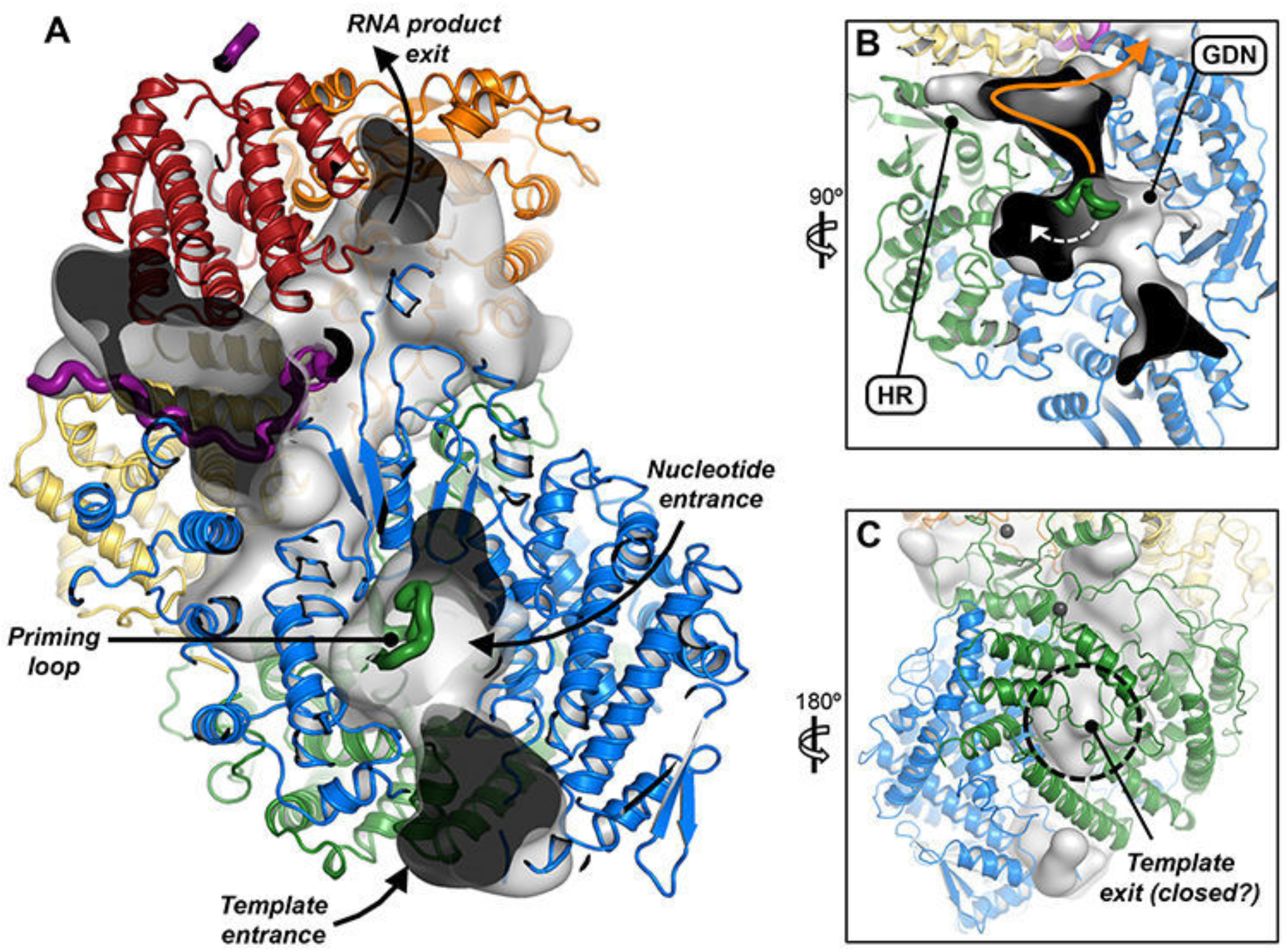
Proposed RNA transit through L. (A-C) Cutaways showing continuous internal cavities (gray, translucent surface) through L (colored, as in Fig. 2C) when the priming loop residues (green ribbon) are deleted from the model. (B) Cavity into which the priming loop may retract during elongation (white dashed arrow). Nascent RNA may continue (orange arrow) toward the CAP active site (HR motif) and MT domain. Black arrows: nucleotide and template entrance channels and the presumed RNA product exit channel. (C) The presumed template exit channel at the back of the CAP domain (black dashed circle) appears “closed” by small loops.

The region with weakest conservation, both of sequence and of structure, surrounds the template entrance channel, the position at which template must separate from N and thread through the catalytic site. Evidence that virus-specific features of L may facilitate this process comes from the observation that RABV L cannot transcribe RNAs from the VSV nucleocapsid, even when the RNP-binding region of P is replaced by that of VSV (Morin et al., 2017). The N-terminal 28 residues of RABV L are disordered, as are the initial 34 residues of VSV L. Their adjacency to the template entrance channel suggests a role in nucleocapsid engagement and template insertion; the low isoelectric point of the disordered peptide further suggests that it might facilitate separation of the template RNA from N.

### Capping domain

The RABV L capping enzyme catalyzes the formation of GTP-capped pre-mRNA by a mechanism that differs from eukaryotic capping. The nascent RNA transcript with a 5’ triphosphate is first covalently linked to a catalytic histidine residue (H1241) in a reaction that leads to a monophosphate RNA-L intermediate. This linkage is subsequently thought to be attacked by a GTP molecule, resulting in addition of GDP to the monophosphate-RNA to form a GTP-capped pre-mRNA (Ogino and Green, 2019). Several residues have been identified in both RABV and VSV CAP that are essential for cap addition, including GxxT and HR motifs broadly conserved throughout *mononegavirales* (Liang et al., 2015). The respective CAP domains are extremely similar (Figs. S5 and S12), although there are small differences in the positioning of certain loops, including those in or near the active site (Fig. S12A). The two sites that coordinate structural zinc ions in VSV L are also present in RABV L (Fig. S12B). Each site includes two pairs of residues separated by approximately two hundred residues in the polypeptide chain, thereby joining separate regions within the CAP domain.

The GxxT motif leads into the ∼20-residue long priming loop, which extends from CAP toward the RdRp active site (Fig. S12C), as we also found for VSV L (Liang et al., 2015). Relatively poor density in both maps indicates conformational variability. The state captured in this structure is evidently a pre-initiation or initiation conformation, as elongation after priming requires that the priming loop withdraw from the RdRp catalytic cavity to make room for nascent product to reach the CAP active site and extend upward into the MT domain (Fig. 5). The catalytic cavity of CAP appears to be large enough to accommodate a retracted priming loop (Fig. 5B), together with a short segment of nascent transcript, even without displacing CAP from RdRp.

### Connector domain

The RABV CD has no known enzymatic function and probably has a largely organizational role. The CD of RABV L consists of eight helices; long linkers at either end connect it to the CAP and MT domains (Fig. S13). Whereas the CAP-CD linker is sufficiently ordered to assign rotamers to several larger side-chains, much of the longer CD-MT linker is disordered, and even main-chain coordinates can be assigned only approximately. Nevertheless, the general course of the latter linker is clear, and we have therefore included the poorly ordered residues in the model (Fig. S13A).

Opposite the HR motif at the CAP active site are three tightly grouped basic resides in the final helix of the CD (Fig. S13B). The first two, Arg1610 and His1611, conserved among all *lyssavirus* L, are also present in VSV L. The third, R1614, is either arginine or lysine in all lyssaviruses; it is a lysine in VSV L. The proximity of these basic residues in the CD to the catalytic HR motif in CAP, as well as their chemical composition, suggest that they may help guide the nascent transcript toward His1241, the position of covalent attachment. Alternatively, they could direct GTP-capped mRNAs into the MT active site.

### Methyltransferase domain

The RABV MT is likely to be a dual-function enzyme that methylates the GTP cap of viral mRNAs, first at the 2’-O and then at the N-7 position, as shown for the VSV MT (Rahmeh et al., 2009). The structurally conserved RABV and VSV MT domains closely resemble other dual-function viral MTases, including those of West Nile Virus (WNV) and Dengue Virus (DENV) (Fig. S14) (Benarroch et al., 2004; Zhou et al., 2007). A shared GxGxG motif forms the binding site for the methyl donor, S-adenosylmethionine (SAM), and a conserved set of charged residues (K-D-K-E; RABV residues Lys1685, Asp1797, Lys1829, Glu1867) forms the catalytic tetrad for methyl group addition (Li et al., 2007; Li et al., 2005; Li et al., 2009; Li et al., 2006). The K-D-K-E tetrads in RABV and VSV MTs are slightly further from the SAM binding site than in WNV and DENV MTs, possibly due to the presence of bound S-adenosylhomocysteine (SAH, modeled) in the WNV and DENV MT crystals; there is no such density in the RABV and VSV L cryo-EM maps.

In addition to the K-D-K-E and GxGxG motifs, we identify neighboring residues in RABV L that are interspersed with the K-D-K-E residues and are close to the presumptive cap binding site (indicated by the location of ribavirin in the DENV MTase structure (Benarroch et al., 2004)): R1674, Y1831, T1860, and Y1869 (Fig. S14B). These residues balance charges of the catalytic residues and may support cap positioning for methylation and/or reaction order. They are conserved throughout *rhabdoviridae* and match similarly positioned residues in WNV and DENV.

### C-terminal domain

The CTDs of RABV and VSV L are the least conserved of the five domains; only 20.1% of the amino acid residues at corresponding positions are the same. Nonetheless, direct alignment of the CTDs shows they are close structural homologs (Fig. 3B), with an rmsd of 3.7 Å.

Density for the CTD was less well defined than it was for most of the rest of the molecule. Multi-body refinement of an expanded particle dataset showed that both the CTD and the MT explore a range of conformations, flexing with respect to the other domains (Fig. S4). While the MT flexes as a rigid domain, the CTD appears to have “open” and “closed” conformations that would involve remodeling the α helix spanning residues 2092-2105 (see the upper left in movies 2 and 3, Fig. S4). This hinge-like behavior may account for the lack of well-ordered density for P at the apex of the CTD in the high-resolution map. Elements of the CTD closest to the MT domain, such as the helix spanning residues 2019-33, follow the MT domain away from the rest of the CTD when “open”, consistent with the close contact between the two sequential domains. The open state in our particle set could represent a fluctuation toward the fully open conformational ensemble seen in the absence of P, in which the CD, MT and CTD all move away from the RdRp-CAP module. We suggest below that capping and methylation may require opening of L in this way.

The C-terminal tail that anchors the CTD in VSV L is absent in RABV L, which has 24 fewer residues. Whereas the C-terminal tail in VSV L-P projects into a cavity in the core of the complex bounded by each of the five domains (Fig. 3B,C), the C terminus of RABV L terminates at the surface of the globular CTD (Fig. 3B). A segment of P that penetrates the gap between CTD and CD, and somewhat closer packing of the adjacent domains, together appear to compensate for the “missing” C-terminal arm (Fig. 3C).

### Priming, initiation, elongation and capping

The structure described here, together with those of VSV L-P at 3.0 Å resolution (Jenni, et al., submitted) and of two recently published pneumovirus L-P complexes ((Gilman et al., 2019) and Pan et al., submitted), suggests a mechanism for switching between replication and transcription, with alternative priming configurations and alternative sites for product exit.

The “priming loop” (residues 1170 to 1186 in the CAP domain) projects into the RdRp catalytic cavity, closing off a channel that connects it with the catalytic cavity of the CAP domain. Model building in the homologous VSV L-P complex shows that an initiating nucleotide would stack on a tryptophan (W1167), corresponding to residue W1180 in RABV L, at the tip of the projecting loop (Jenni et al, submitted). Mutation of this residue compromises end initiation, but not internal initiation or capping (Ogino et al., 2019). Elongation beyond formation of the initial dinucleotide requires that the priming loop retract into the CAP catalytic cavity. The recent pneumovirus L-P structures show just such a retracted configuration (Gilman et al., 2019) and Pan et al., submitted. A nascent transcript can then pass across the retracted loop. Moreover, analysis of cavities in the L-P complex (Fig. 5) shows that after priming loop retraction, a continuous tunnel leads from the connected catalytic cavities of RdRp and CAP to a likely exit site for the full-length replication products (antigenomic and genomic RNA). Proximity of this putative exit site to the N-terminal end of P would then allow prompt delivery of N protein, bound near the N-terminus of P, coating and protecting the emerging replication product (or uncapped leader transcript).

The N-protein delivery pathway just suggested depends on the site at the apex of the CTD that anchors the N-terminal segment of P. That site is a shallow pocket, which accommodates just four or five residues. In the VSV L-P complex, a tyrosine side chain packs against the base of the pocket; mutation of that tyrosine to small residues severely impairs viral growth, but does not affect RdRp activity (Jenni et al., submitted). We have suggested that failure to anchor P in that site may impede replication (but not transcription) by interfering with efficient delivery of N to the product. That function might explain the conservation of the position of the site, but not its chemistry.

Priming for internal initiation and capping and priming for end initiation may depend on different configurations and different mechanisms for supporting the priming GTP. Internal initiation is insensitive to mutation of Trp1180 (Ogino et al., 2019), suggesting that the priming loop remains retracted after termination of the leader transcript and that some other structure – perhaps the 3’ end of the preceding transcript, still based paired with template – provides support for the priming GTP (see a related, early proposal in (Shuman, 1997)). Each of the products of internal initiation, with a 5’-AACA sequence, forms a covalent attachment with His1241. A noteworthy characteristic of the “closed”, initiation competent structure seen here and for VSV L is the absence of an obvious GTP binding site in the capping domain. We suggest that the retracted priming loop and elongated nascent transcript might create such a site, perhaps linked to covalent attachment of the 5’ end of the transcript to His1241. As long as the 5’ attachment to His1241 is present, elongation will produce a loop that will fill the CAP catalytic cavity and force the smaller domains (CD, MT, CTD) to swing outwards – as they do in the absence of P. This transition to a more “open” complex will allow GTP to diffuse into the CAP active-site cavity (if it is not already there), and it will also expose the methyltransferase catalytic site more completely than in the closed structure. These steps may account for the observation that capping of VSV mRNA occurs only after 31 nucleotides have been transcribed (Tekes et al., 2011)

The scheme proposed here provides a simple mechanism for switching between replication mode, in which full-length product acquires an N-protein coat, and transcription mode, in which an mRNA product acquires a 5’ cap. It also provides an evolutionary rationale for the distinctive capping mechanism (polyribonucleotide transferase rather than guanylyl transferase) found in NNS viruses. Retraction of the priming loop creates a continuous cavity shared by the CAP and RdRp catalytic sites. Covalent attachment of the 5’ end of the transcript then allows on-going RNA polymerization to fill this cavity with product strand, generating the force needed to release the CD-MT-CTD module. Tests of this model will require structures of transcribing intermediates.

## ACKNOWLEDGMENTS

We thank Louis-Marie Bloyet for discussions of mononegavirus biology and advice on L-protein preparation. JAH is an Amgen Fellow of the Life Sciences Research Foundation. The work was supported by NIH grants R37 AI059371 (to SPJW) and R01 CA13202 (to SCH). SCH is an Investigator in the Howard Hughes Medical Institute.

## AUTHOR CONTRIBUTIONS

All authors designed research; J.A.H. performed research; J.A.H. and S.C.H. wrote the paper; all authors edited the paper.

## METHODS

### Protein expression and purification

We expressed 6xHis-tagged recombinant RABV SAD-B19 L alone or with variants of RABV SAD-B19 P in SF9 or SF21 insect cells from baculovirus vectors constructed using the pFastBac-Dual recombination system (Thermo Fisher) as described previously (Morin et al., 2017). The N-terminal fragments of RABV SAD-B19 P, P_11-50_ and P_1-91_, were expressed in *E. coli* and purified as previously described (Morin et al., 2017). The P variants GFP-P_1-91_, P_FL_, and P_ΔOD_, with StrepII affinity tags and were co-expressed with L in insect cells and co-purified with L. For GFP-P_1-91_, with an N-terminal StrepII tag, L-GFP-P_1-91_ complexes were purified first on nickel and then on Strep-Tactin resin (GE Healthcare). For P_FL_, with a C-terminal StrepII tag, L-P_FL_ complexes were purified first on Strep-Tactin and then on nickel resin. For P_ΔOD_, with an HA epitope tag followed by an internal HRV 3C protease cleavage site in place of the deleted oligomerization domain of P (P residues 92-131), as well as a C-terminal StrepII tag, L-P_ΔOD_ complexes were immobilized on Strep-Tactin resin, then cleaved from the resin in the presence of HRV 3C protease to yield purified complexes of L-P_1-91_. The 9-residue HA tag, not used for purification, remained attached to P_1-91_ following proteolytic cleavage. For cryo-EM specimens, these L-P_ΔOD_-derived L-P_1-91_ complexes were further purified on a Superdex 200 Increase column and the peak fractions concentrated to 0.3 mg/ml immediately before grid preparation. For each grid, 3ul of sample were applied to glow-discharged, C-Flat 400-mesh copper grids coated with 40 nm thick holey carbon (1.2/1.3 spacing) and plunge-frozen in liquid ethane in a CP3 cryo-dipper.

### Electron microscopy

For the medium-resolution initial reconstruction, images of RABV L-P_1-91_ were recorded on a Tecnai F20 electron microscope (FEI) operated at 200 kV, using UCSF Image4 (Yuemin Li, UCSF) to collect movies on a K2 Summit direct detector (Gatan) in superresolution mode with dose fractionation. For each 8-s exposure, we collected 32 frames at 250 ms each, with a total electron dose of 72 e/Å^2^. Collection was performed at a nominal magnification of 29,000X, with a calibrated pixel size of 1.28 Å/pixel. Movie frames were gain-subtracted, Fourier-binned 2X, and motion-corrected using MotionCor2 with 5×5 patch correction.

For the high-resolution dataset, images were recorded on a Tecnai F30 Polara electron microscope (FEI) operated at 300 kV, using SerialEM to collect movies on a K2 Summit direct detector (Gatan) in superresolution mode with dose fractionation. For each 8-s exposure, we collected 32 frames at 250ms each, with a total electron dose of 72 e/Å^2^. Collection was performed at a nominal magnification of 31,000X, with a calibrated physical pixel size of 1.234 Å/pixel.

### Image processing

Micrograph movie frames were gain-subtracted, Fourier-binned to physical pixel size, and motion-corrected using MotionCor2 with 5×5 patch correction. Micrograph CTF coefficients were calculated using CTFFIND4 (Rohou and Grigorieff, 2015) and particle images were CTF-corrected as needed by the processing software used.

For the medium-resolution reconstruction, a single dataset was collected from one of four identical grids prepared in parallel from a single protein sample. We picked 43,000 particles by hand from 316 motion-corrected movies using EMAN2.1(Ludtke, 2016). A de novo 3D reference was obtained using the ab-initio procedure in cisTEM(Grant et al., 2018) following preliminary 2D classification, and the hand subsequently flipped to yield a reference bearing strong resemblance to VSV L-P. Final 2D and 3D classification using the flipped reference were then carried out in RELION 2.1(Kimanius et al., 2016), yielding a 6.7Å (medium-resolution) map from 23,000 particles.

For the high-resolution reconstruction, we collected a total of 10,949 movies from three datasets using the remaining three grids from the initial set of four. Selected 20 Å-low-pass-filtered projections of the 6.7 Å map were used as particle-picking templates for autopicking in RELION. Autopicked particles on each micrograph were manually cleaned of junk, contamination, noise, drift, and otherwise bad or damaged particles using SamViewer, an interactive image analysis program written in wxPython by Dr. Maofu Liao (Harvard Medical School). From 5.3 M remaining particles, coarse 2D classification at a high-resolution cutoff of 25 Å was performed in RELION to further remove poor particles, and classes representing dominant views were separated from the remaining good classes, as described in Fig. S2. 3D classification was performed on 1.47 M non-dominant view particles, and the best class of six was selected, containing 649 k particles. These particles were added to the separated 408 k dominant view particles; a final round of 3D classification then yielded a total of 963 k particles from the best class of three. This particle set was then subjected to 3D refinement, per-particle CTF-refinement, and Bayesian polishing in RELION 3.0, and the resulting stack imported into cisTEM along with corresponding Euler angles and offsets for further refinement, as described in Figs. S1 and S2. The anisotropic 3D reconstruction from refinement in RELION was used as a starting reference for manual refinement in cisTEM, and particles were removed from the reconstruction at each refinement iteration using the SCORE parameter. We quickly removed the worst-scoring 50% of particles. Because dominant view particles had higher scores (due to anisotropy of the map caused by a preferred orientation), we applied an angular binning script to cap the number of particles per 4×4-degree bin, keeping those with the highest scores. Further manual refinement using these remaining 177 k particles with flattened angular distribution led to reduced anisotropy and improved map quality, with the best 25% of particles by score giving the highest-resolution map. A final refinement using this set of 44,500 particles at an alignment resolution cutoff of 4.5 Å yielded a reasonably isotropic map (despite a majority of particles having the preferred orientation) with an average resolution of 3.3 Å.

### Model building

To generate the atomic model of RABV L, we first replaced the sequence of VSV L with RABV L in the model of VSV L (Jenni et al., submitted) adding simple loops where discontinuities in the alignment occurred. We fit these coordinates as a single rigid body into the 3.3 Å RABV L-P_1-91_ density map, then divided the coordinates into the five structural domains (RdRp, CAP, CD, MT, and CTD) and re-fit each domain as a rigid body. We then performed iterative real-space refinement using PHENIX (Adams et al., 2010; Afonine et al., 2018) at increasing resolution, from 40 Å to 5 Å. The resulting model was then used as an input model for ROSETTA(Song et al., 2013). To improve loop fitting, we also input libraries of 3-mer and 9-mer peptide structures corresponding to all 3- and 9-amino acid sequences contained in the 2127-residue RABV L sequence (Kim et al., 2004). From 5,000 simulations, we selected a model with very good map-to-model agreement and with visibly strong fitting of secondary structure elements throughout. We then performed several rounds of manual adjustment in Coot, alternating with real-space refinement in PHENIX, using secondary structure restraints throughout and Ramachandran restraints in the final rounds. The final model was validated using PHENIX and MolProbity (Table S1).

### Projection matching

Molecules of RABV L in complex with GFP-P_1-91_ were visualized by negative-stain EM at ∼0.01 mg/ml total protein in the presence of 0.7% uranyl formate. 28 2D class averages of picked particles showed clear density for the GFP moiety at the N-terminus of P (Figs. 4A, S6 and S7). We aligned the 2D class averages with obvious GFP density to a single reference to standardize their orientation in the image plane, and applied a circular noise mask to obscure the GFP signal as much as possible without obstructing signal from L. To determine the viewing angle of the RABV L atomic model corresponding to each 2D class, we projected the high-resolution RABV L-P_1-91_ cryo-EM map over all viewing angles, incremented every ∼7.2° for a total of 800 projections. Projections were low-pass filtered to 20 Å, then scaled and clipped to match the pixel and box size of the 2D classes, and a circular noise mask applied to obscure the clipping edges. Cross-correlation coefficients between projections and 2D classes were obtained using e2simmx.py from EMAN2 allowing for rotation and translation only. The top projection match for each 2D class is shown in Figs. 4 and S6 with GFP modeled as a best-approximation.

L-P_FL_ dimers, representing about 25% of L species on micrographs of negatively-stained, purified RABV L/P_FL_, were picked by hand and subjected to 2D class averaging using EMAN2.1. From 29 high-quality 2D class averages of dimers, we extracted each of the 58 dimeric monomer averages as individual particles (hereafter, DMAPs). We applied a circular noise mask to each DMAP to obscure excess signal from the paired DMAP present in each particle image. We then projected the high-resolution RABV L-P_1-91_ cryo-EM map over all viewing angles, incremented every ∼7.2° for a total of 800 projections. Projections were low-pass filtered to 20 Å, then clipped and scaled to match the box and pixel size of the nsEM DMAPs, and a circular noise mask applied to obscure the clipping edges. Cross-correlation coefficients between projections and DMAPs were obtained using e2simmx.py allowing for rotation and translation only. The resulting matrix was then normalized within each DMAP over all projections, from which we constructed spherical heat-maps in the coordinate system of the RABV L-P_1-91_ cryo-EM map to facilitate visualization of peaks reflecting high cross-correlation scores. For about half of the DMAPs, a single peak predominated, and application of the angular coordinates from the peak to the atomic model of RABV L-P_1-91_ returned an obvious match to the corresponding DMAP. When more than one high-scoring peak was observed, the best apparent visual match was selected. Of the 58 DMAP:projection comparisons, only 6 were matched to a secondary peak, and 5 could not be reliably matched. In total, we successfully matched 24 of the 29 DMAP pairs. Using PyMOL, we then applied the angular coordinates from each match to an atomic model of RABV L-P_1-91_ for each DMAP in a pair, and approximated the relationship between the two modeled DMAPs using only two dimensions (X and Y), as suggested by the original 2D class average. We then measured the minimum distance between C-alpha atoms of either P_87_ or the mapped, but unassigned, N-terminal-most residue of P in each modeled pair by translating one of the models along the Z axis. The results of these projection matching analyses are summarized in Fig. 4B and illustrated in detail in Fig. S8.

### Visualization of internal cavities

We used the program VOIDOO (Kleywegt and Jones, 1994) to probe internal cavities, substrate entry and product exit channels of the RABV L-P_1-91_ complex. We used a probe radius of 1.8 Å to calculate a probe-occupied volume.

## SUPPLEMENTAL INFORMATION

**Figure S1:**
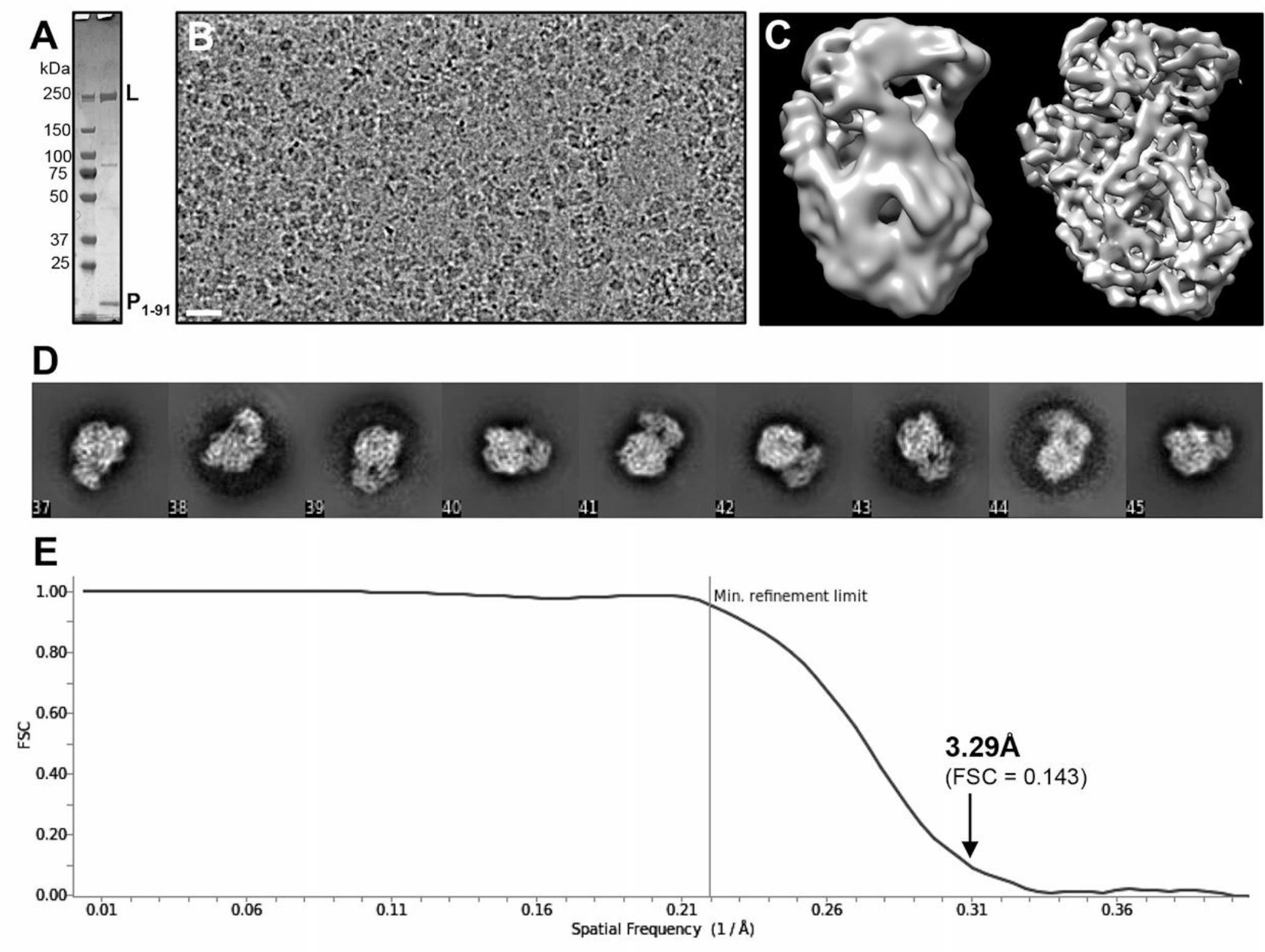
Cryo-EM reconstruction. (A) Coomassie-stained SDS-PAGE gel showing the purified RABV L-P_1-91_ preparation used for cryo-EM. (B) Representative cryo-EM micrograph of RABV L-P_1-91_ complexes from (A) immobilized in vitreous ice and imaged at 300 kV. Scale bar: 25nm. (C) Ab-initio 3D reference (left) and 6.7 Å reconstruction (right) from a preliminary 200 kV dataset. (D) Representative 2D class averages from the final 44.5K particle dataset (Fig. S2) used to obtain the 3.3 Å reconstruction. (E) Fourier shell correlation curve for the final 3.3 Å map (cisTEM).

**Figure S2:**
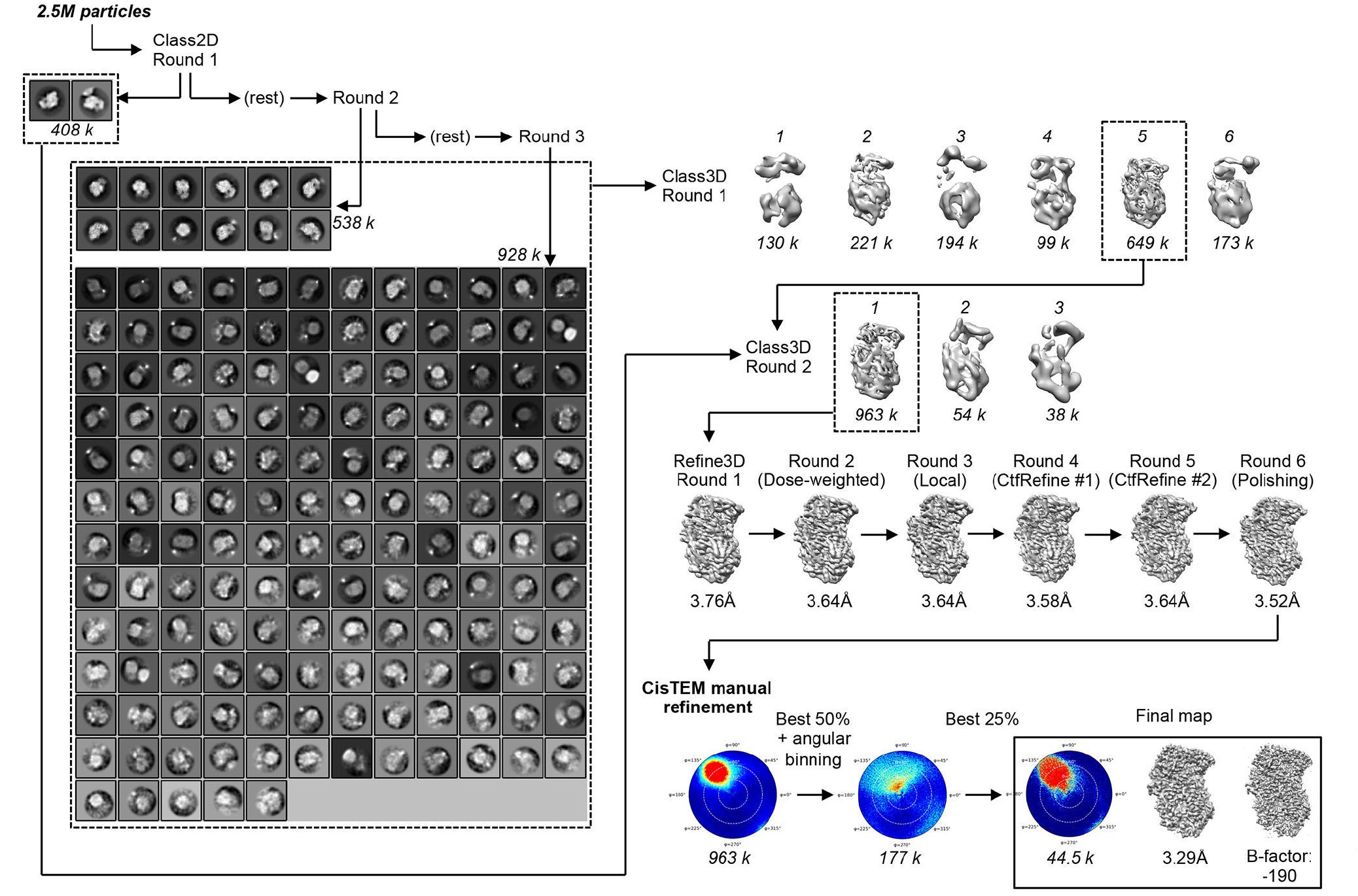
Particle classification. From 5.3 M initial particles after auto-picking and manual cleaning, particle images from each of the three 300 kV datasets were binned 3x and subjected to an initial round of 2D classification in RELION to remove bad particles (not shown). The remaining 2.5 M particles were subjected to a round of coarse (25 Å resolution cutoff) 2D classification (Class2D Round 1). The two classes with the highest occupancy, containing a large portion of the particles adopting the preferred orientation we observed, were separated from the remaining particles and retained for later use. In a second round of 2D classification with the remaining particles, 12 good classes totaling 538 k particles were selected and again separated from the rest, to be used later. The remaining particles were subjected to a third round of 2D classification, from which all but the very worst classes were kept to maximize retention of particles with rare views, totaling 928 k particles. These particles were pooled with the 538 k particles selected in round 2, and the combined 1.47 M particles subjected to 3D classification (Class3D Round 1). The best class among six was chosen for further classification, containing 649 k particles. These particles were merged with the additional 408 k preferred-orientation particles separated in the first round of 2D classification, and the combined 1.06 M particles subjected to a round of 3D classification (Class3D Round 2). A single class predominated, with 963 k particles. We extracted unbinned particles images of this class for several rounds of 3D refinement using automated Refine3D in RELION. After an initial round using non-dose-weighted particles, subsequent rounds used dose-weighted particles, iterative per-particle CTF-refinement, and Bayesian polishing. Although modest gains in reported resolution were observed, the map remained visibly anisotropic even after particle polishing. The entire polished stack of 963 k particles was then imported into cisTEM, and several rounds of manual refinement were carried out in which particles were removed on the basis of the ‘SCORE’ parameter and by angular flattening. Lower right: Heat-maps (cisTEM), showing the distribution of Euler angles at the outset and after initial and final particle pruning, and two representations of the final map, with an average resolution of 3.29 Å (FSC = 0.143 criterion), reconstructed from 44,500 particles.

**Figure S3:**
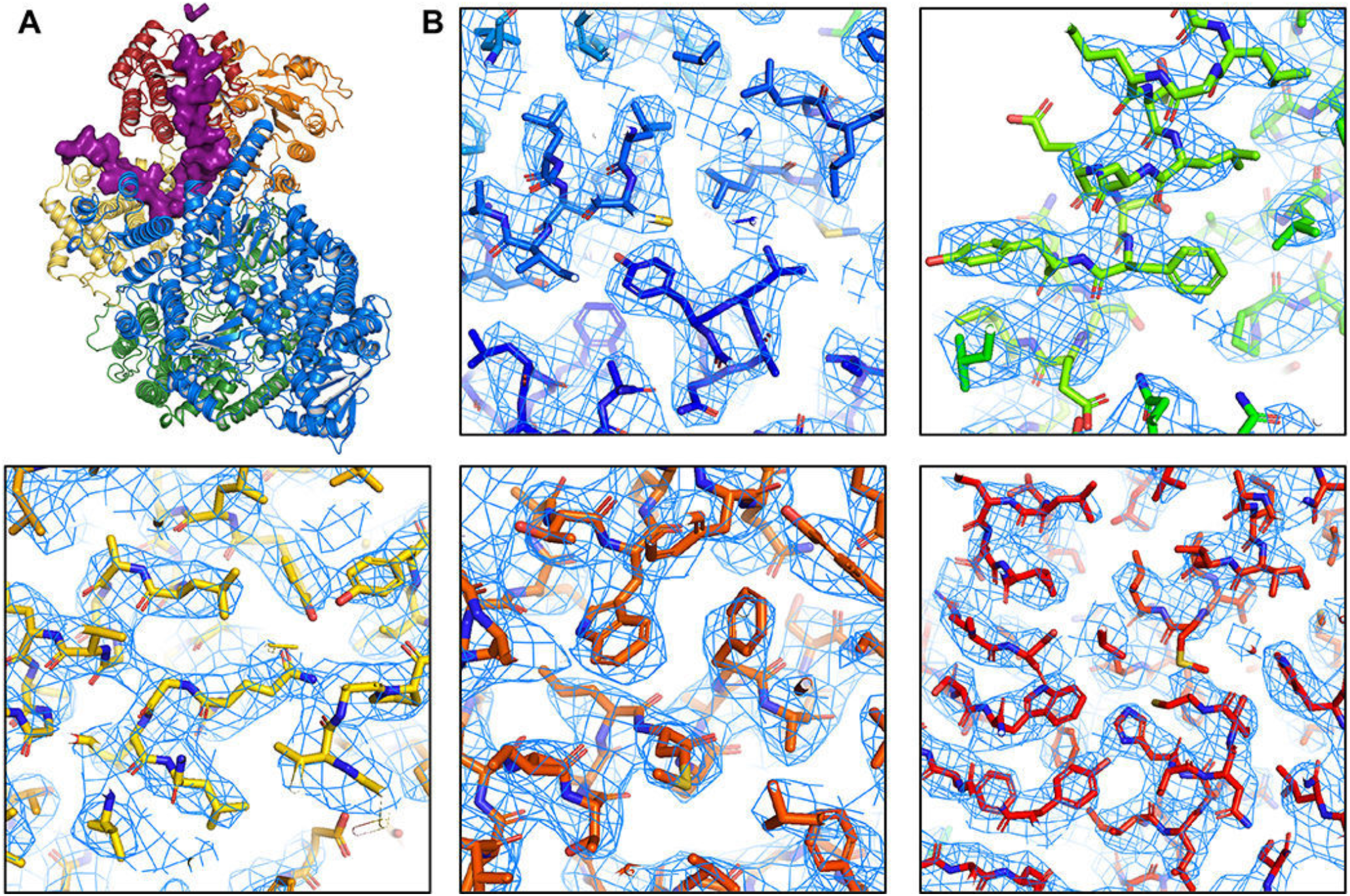
Details of 3.3 Å map. (A) RABV L-P_1-91_ structure, as in Fig. 2. (B) Side-chain density in each of the five domains of L. Atomic model is in stick representation. Carbon atoms colored by domain, as in (A) and Fig. 2; oxygen, red; nitrogen, blue; sulfur, yellow.

**Figure S4:**
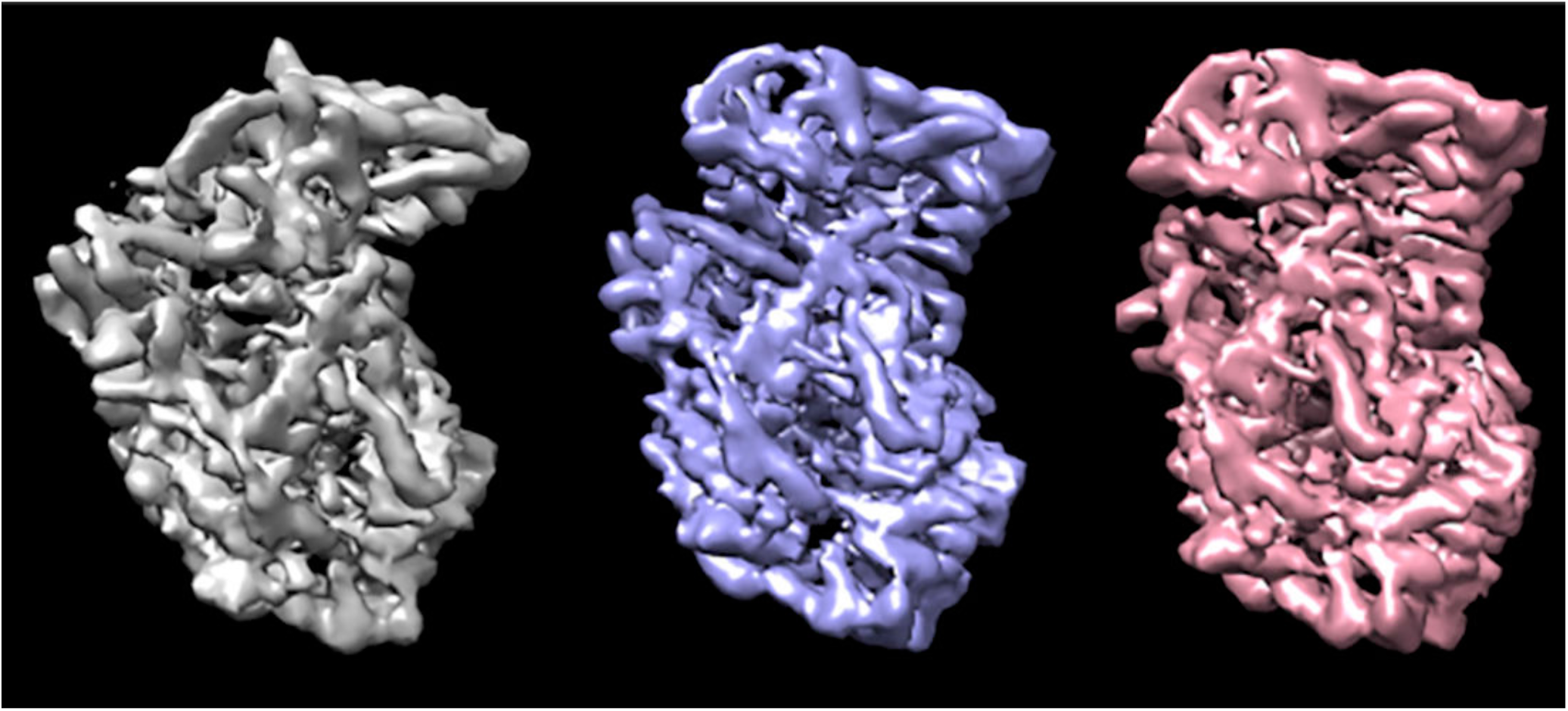
RABV L conformational variability. Multi-body refinement in RELION 3.0 with four masked regions (RdRp+CAP; CD; MT; CTD) using an expanded dataset of 2.4 M particles shows a broad range of conformations for the C-terminal domains of RABV L. “Movies” illustrate conformations explored along the three eigenvectors (of 24 calculated) accounting for the greatest variance in the data. Each movie has ten frames, with each frame a reconstruction from 1/10^th^ of the particles populating the given eigenvector. The frames in each movie were aligned to the RdRp domain of RABV L to illustrate the different positions of the C-terminal domains (CD, MT, CTD) with respect to each other and to the RdRp-CAP domains. The movies are not aligned to one another; each is oriented to show most clearly the conformations for the corresponding eigenvector.

**Figure S5:**
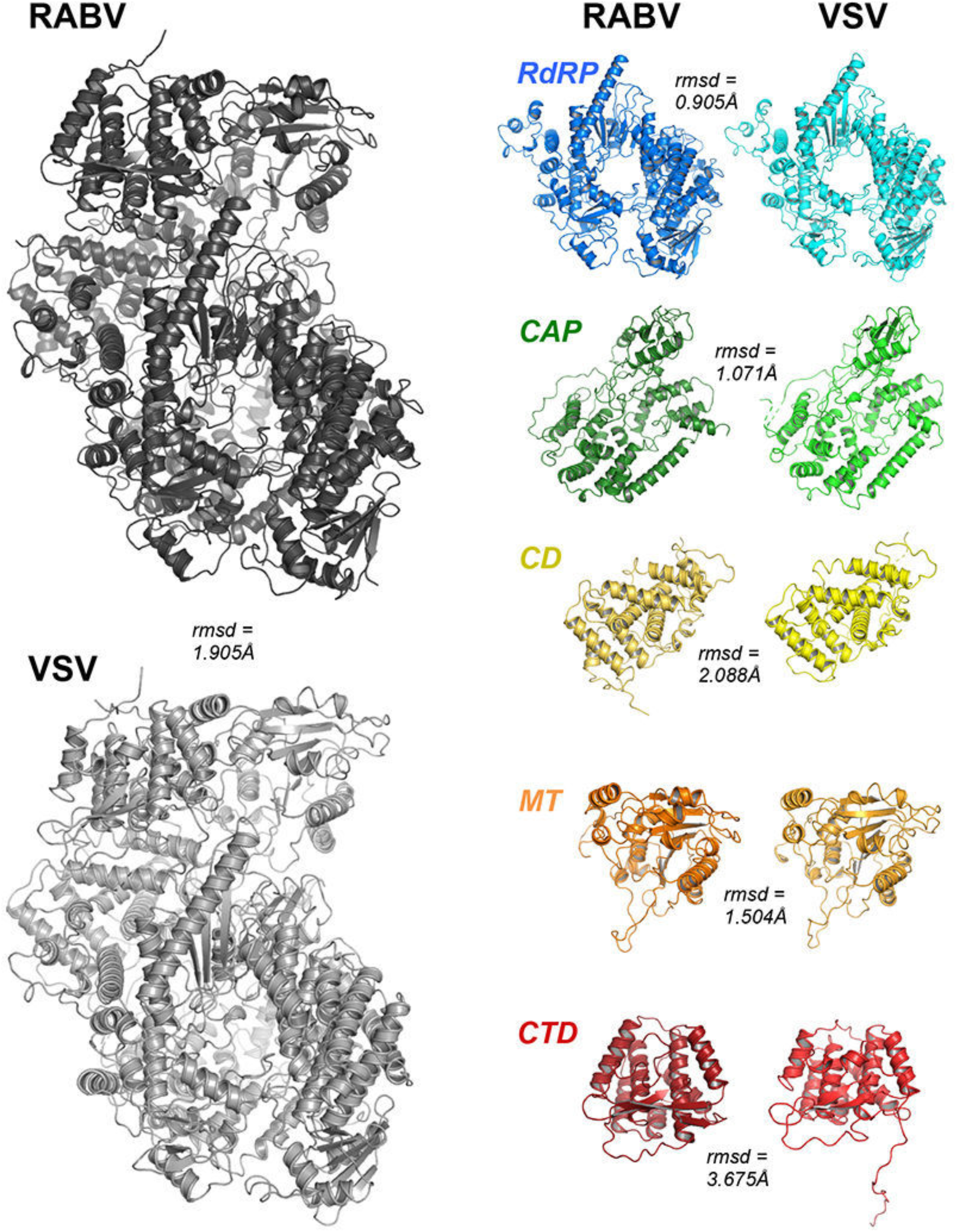
Alignments of RABV and VSV L and their domains. *Left:* RABV (dark gray) and VSV (light gray) L-P structures in cartoon representation with the RMSD after iterative whole-structure alignment of Cα atoms with outlier elimination in PyMOL. *Right:* independently aligned domains of RABV and VSV L in cartoon representation (colored, as in Fig. 2C) with their respective RMSDs.

**Figure S6:**
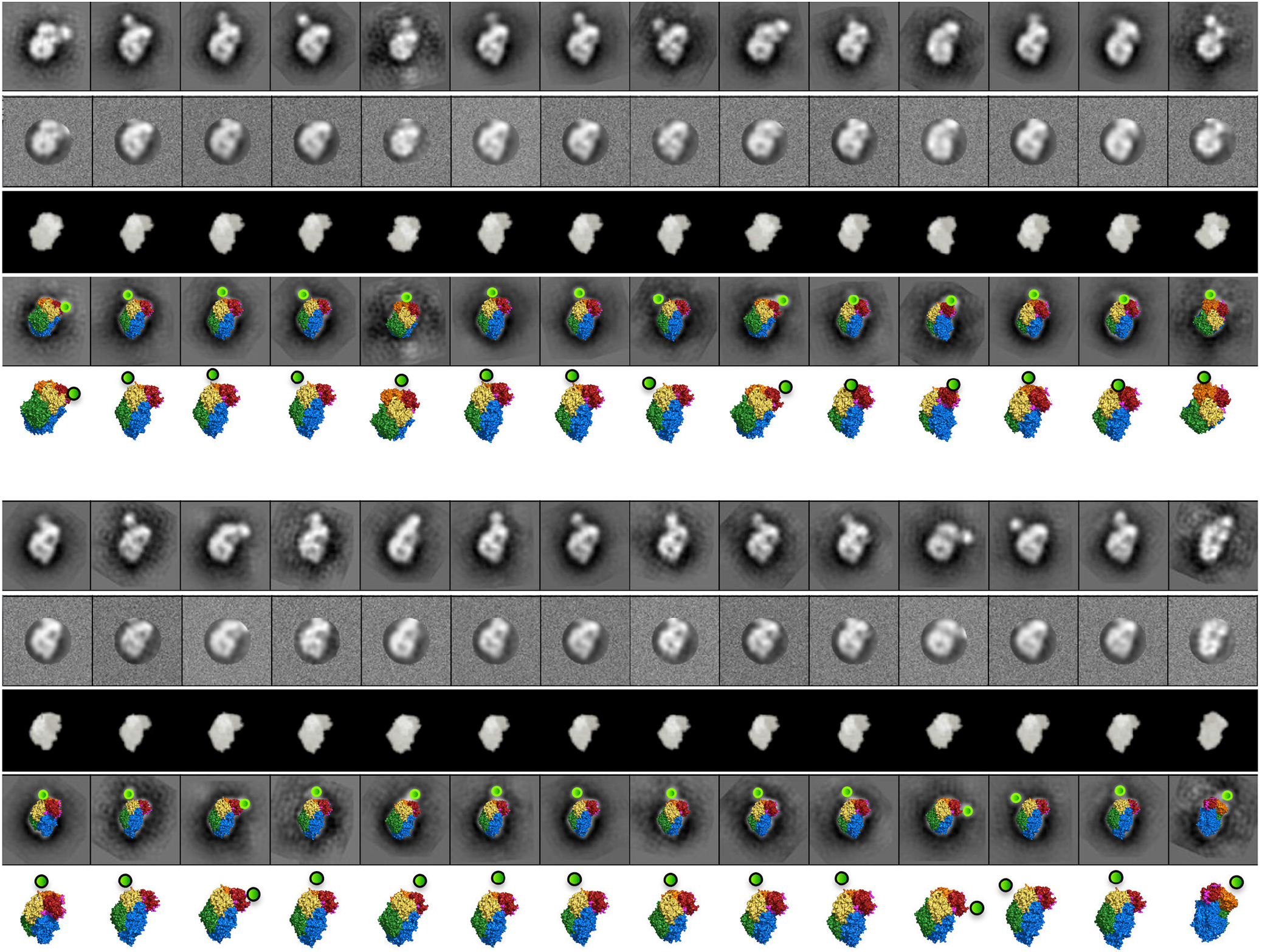
L-GFP-P_1-91_ projection matching. Twenty-eight negative-stain EM 2D class averages of RABV L in complex with GFP-P_1-91_ for which a clear feature corresponded to the globular GFP moiety, after alignment to a 2D reference to standardize their orientation in the X/Y plane (top rows). The 2D class images were noise-masked to remove as much of the GFP signal as possible without losing information from L (second rows). Corresponding matched projections of RABV L were obtained as described in Methods (third, fourth and fifth rows). We then made a best approximation for the location of the GFP globule relative to each projection of RABV L (fourth rows) and scaled the resulting models for ease of visualization (fifth rows). The preferred orientation of the particles is evident from the low diversity of projections represented; the preferred view helps visualize the wide arc defining the N terminus of P in RABV L, as the GFP globule occupies many different positions about several near-identical projections of L.

**Figure S7:**
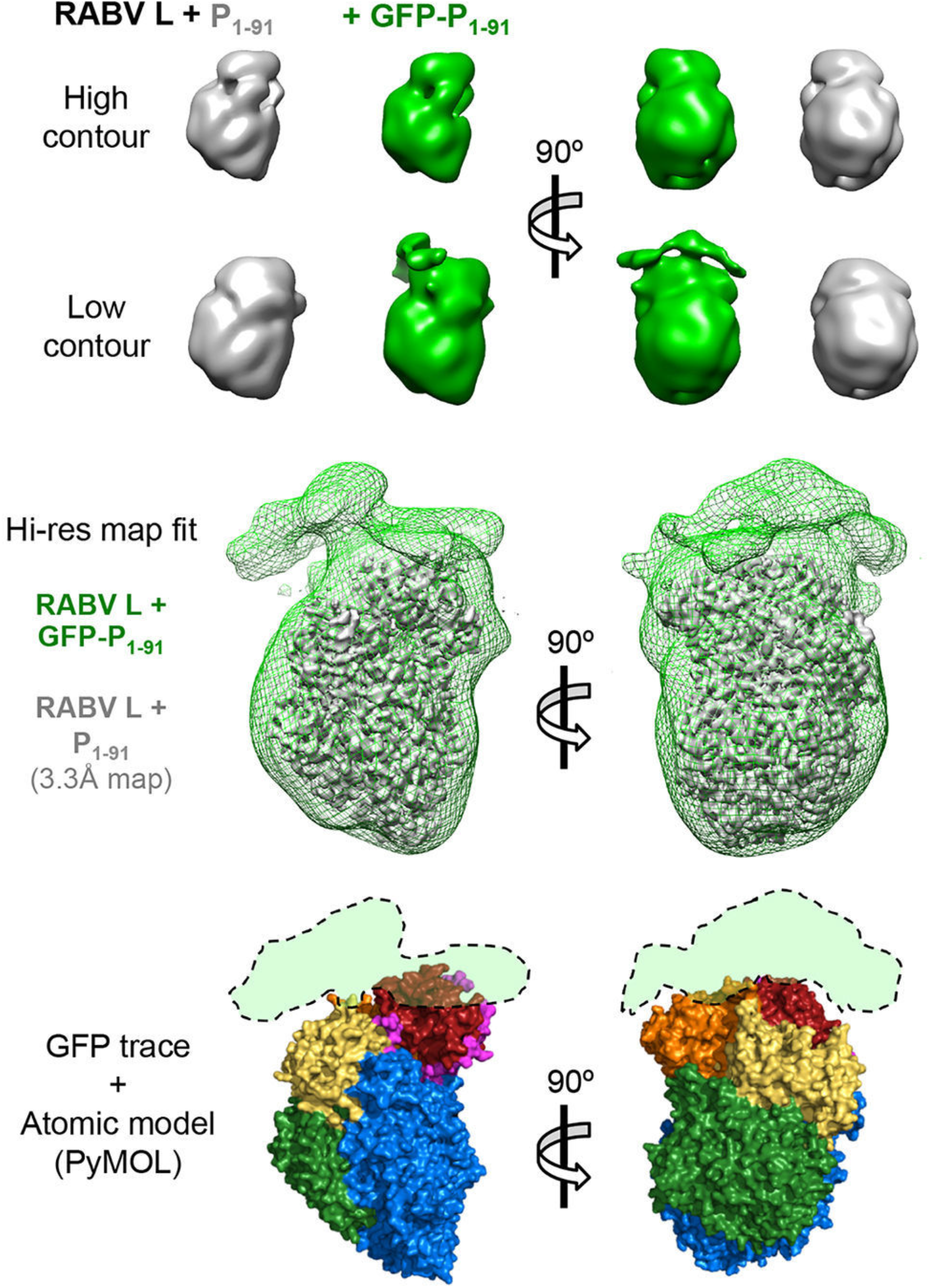
L-GFP-P_1-91_: comparison with high-resolution structure. *Top,* Low-resolution negative-stain EM density maps are shown for RABV L in complex with either P_1-91_ (gray) or GFP-P_1-91_ (green) at high and low contour levels. *Middle*, the high-resolution RABV L-P_1-91_ cryo-EM density map fit (with UCSF Chimera) into the map of negatively stained L-GFP-P_1-91_ complex. The L-GFP-P_1-91_ map is shown at low contour to illustrate the diffuse density for the GFP globule about the top of RABV L. *Bottom*, a crude trace of the GFP density, superposed on the atomic model of RABV L corresponding to the high-resolution map fit above. This image is reproduced in Fig. 4A.

**Figure S8:**
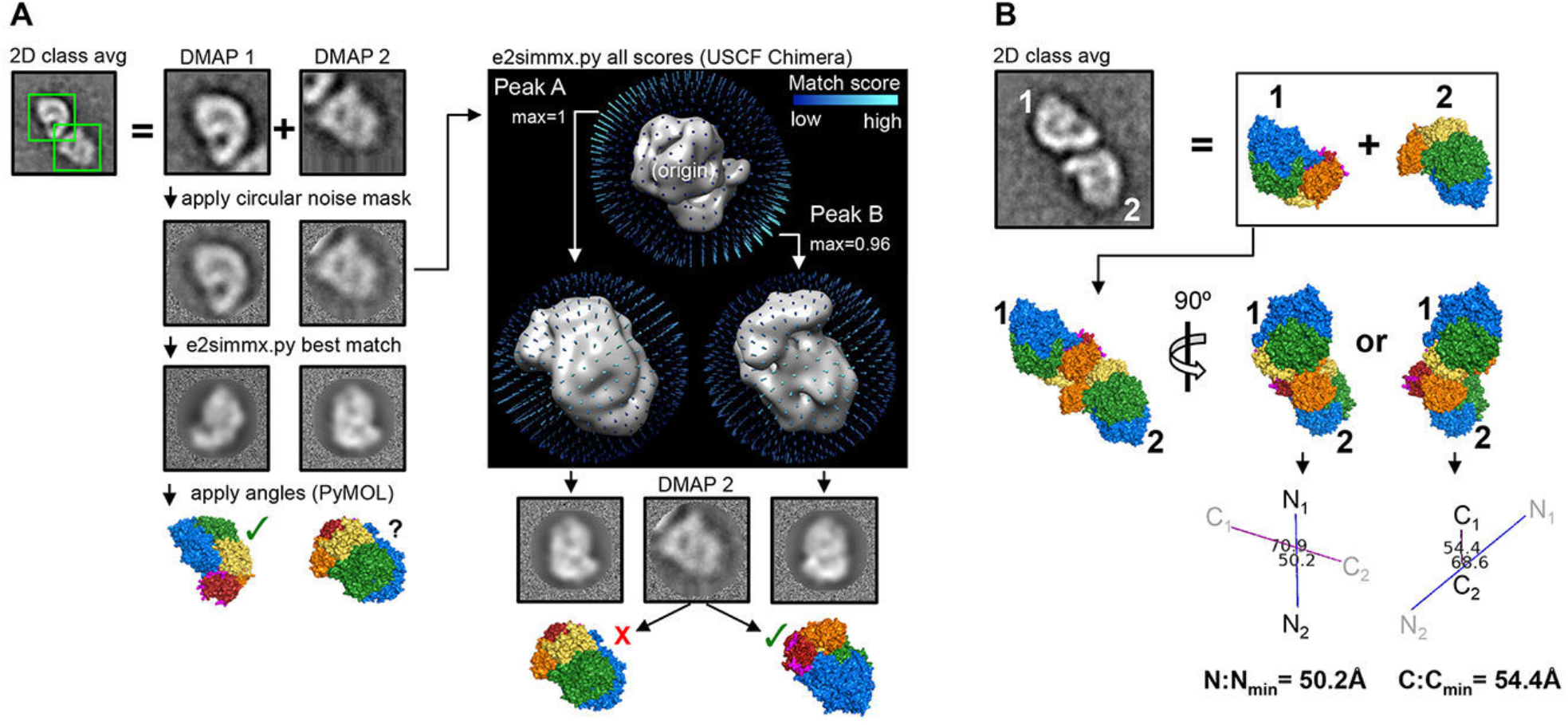
L-P_FL_ projection-matching. (A) Projection matching of L-P_FL_ dimers, as described in Methods, illustrating a case in which one of the partners resulted in an incorrect top projection match. After application of the top projection-matched angles for each dimer partner in PyMOL, the resulting RABV L model images were inspected for correctness. We generated 3-dimensional heat maps for the e2simmx.py projection scores in the coordinate system of the EM density map to visually evaluate the peaks. When obvious, a secondary or alternate peak was selected instead of the top match. (B) Projection-matched models for each monomer (DMAP) in a dimer pair 2D class average were arranged in the X/Y plane in PyMOL to a best-approximation with the original 2D class average. To estimate distances between dimer partners, we translated one of the models along the Z axis such that the distance was minimized between Cα atoms of either the C-terminal or the N-terminal residues of P. This optimization yielded two measurements for each dimer, which we refer to as the C:C_min_ or N:N_min_ distance.

**Figure S9:**
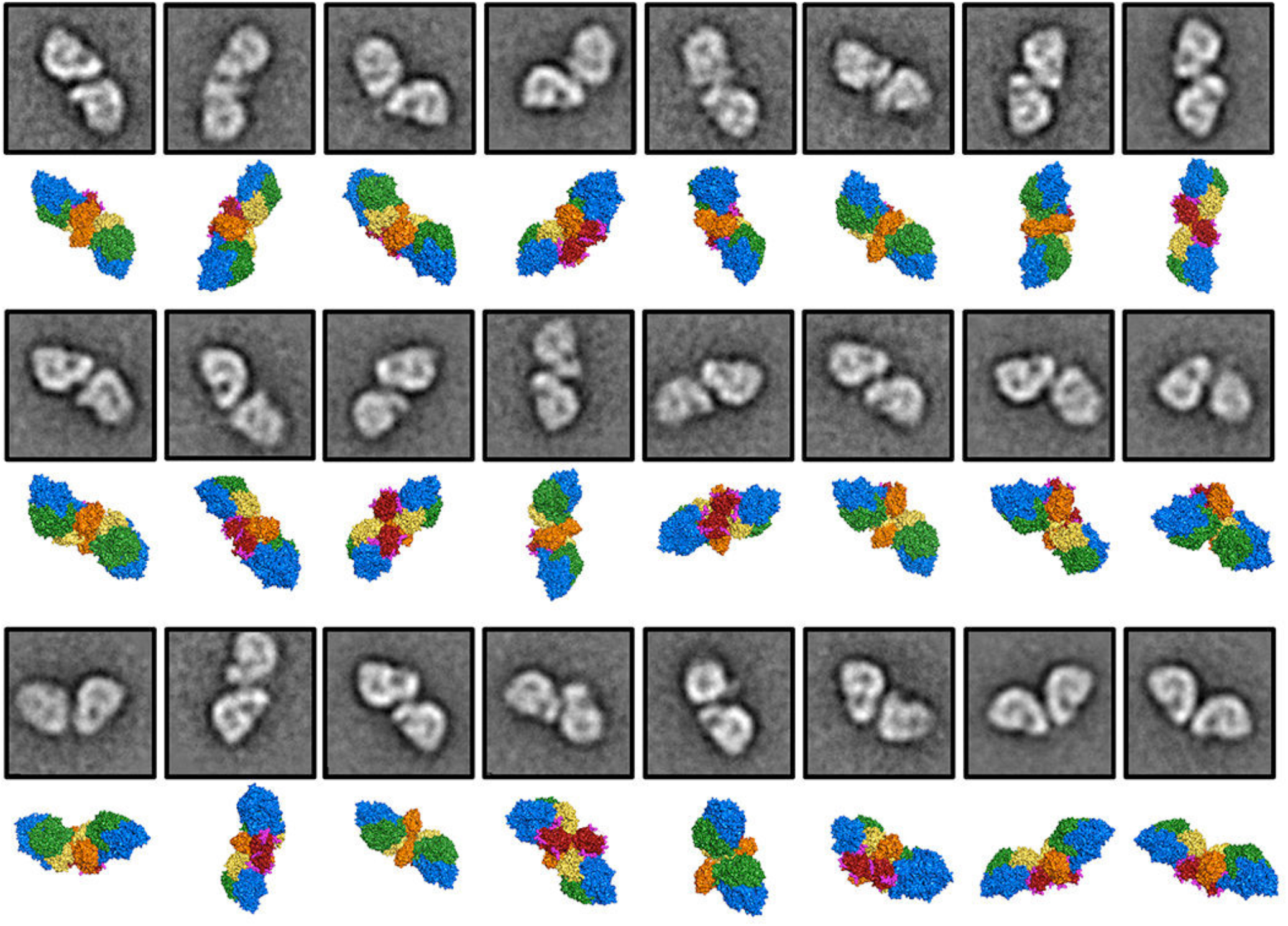
L-P_FL_ projection-matched dimers. Negative-stain EM 2D class averages (top rows) and projection-matched RABV L models (bottom rows) are shown for the 24 dimer pairs (of an initial 29 pairs, not shown) for which we could match both DMAPs.

**Figure S10:**
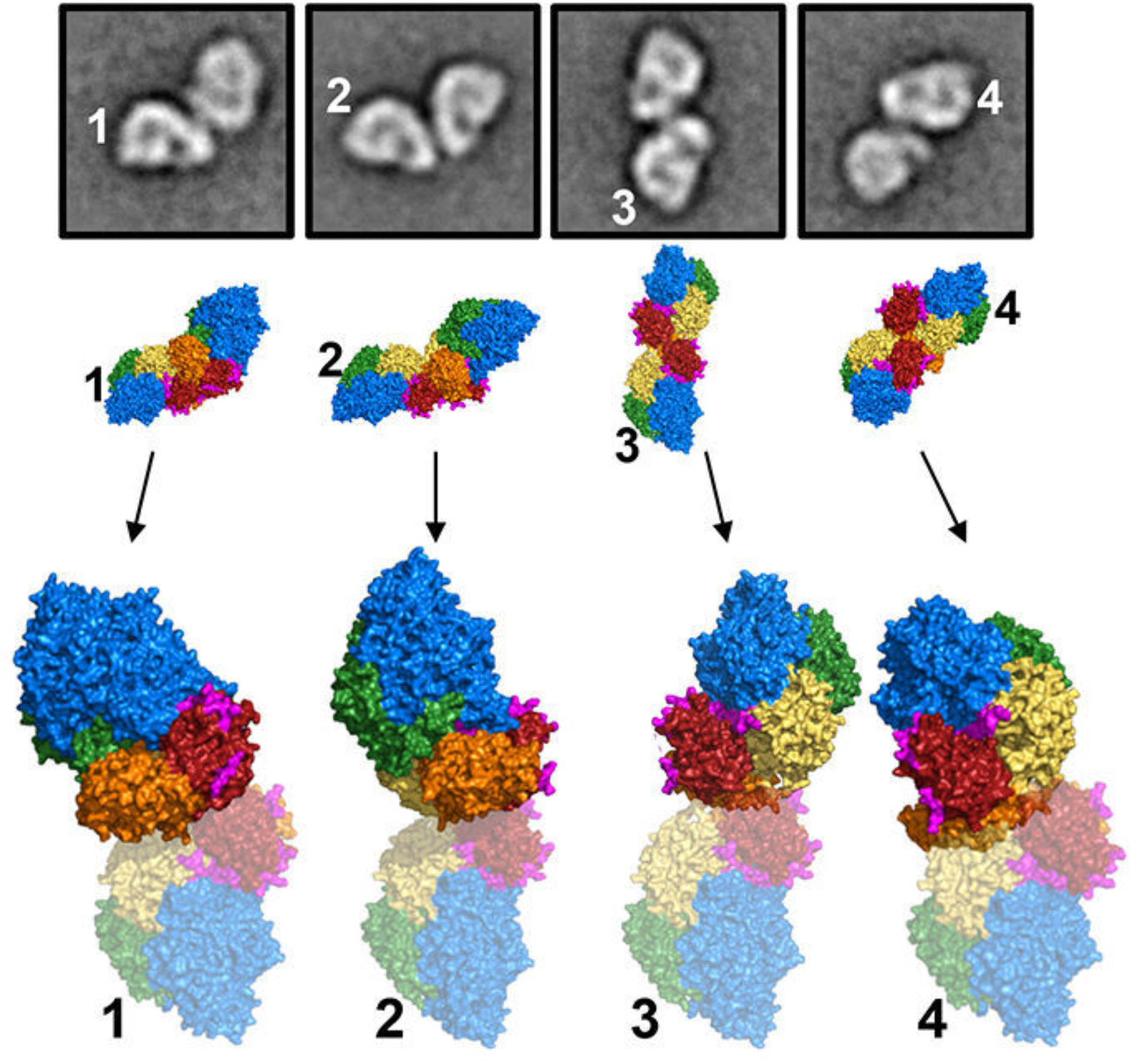
Arrangement of L-P_FL_ dimers. *Top,* Four projection-matched dimers are shown (as in Figs. 4B and S7) for which one of the dimer partners in each shares a common projection with the others. The partners in each dimer sharing the common projection are listed 1-4. *Bottom*, the four projected dimers were rotated about the Z axis such that the common projections were aligned with each other. The common projections are shown partially transparent to show that the partners in each dimer adopt different positions with respect to the common projection.

**Figure S11:**
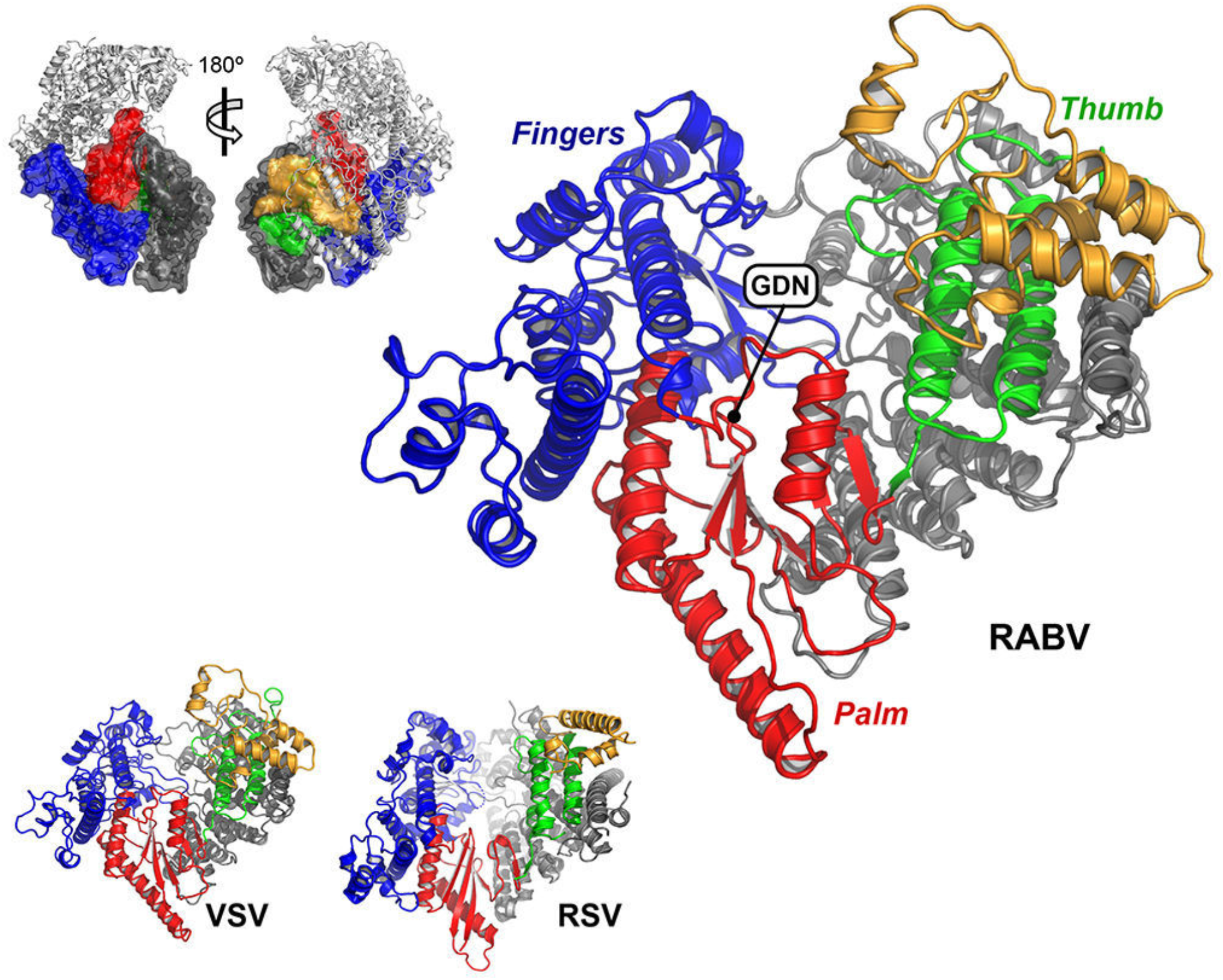
The RdRp. *Right and bottom:* the RABV, VSV and RSV RdRps adopt the familiar “right hand” configuration of nucleotide polymerases, and are shown in cartoon representation: fingers, blue; palm, red; thumb, green; N-terminal extension, gray; and C-terminal extension, orange. *Top-left:* subdomains of the RABV RdRp (surfaces, colored as at right) are highlighted against the complete RABV L-P structure (gray cartoon) to illustrate their positions in L relative to the orientation shown at right.

**Figure S12:**
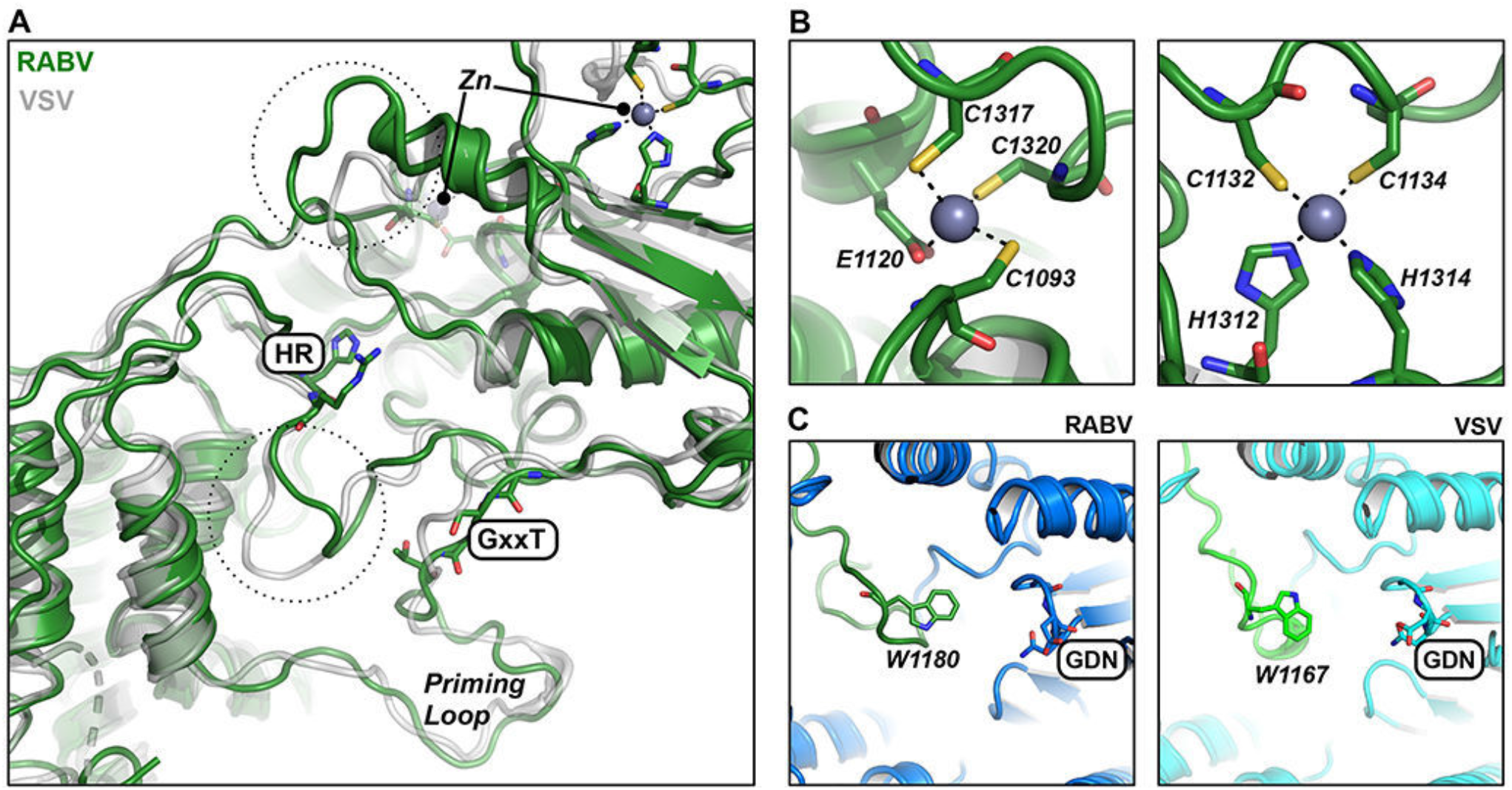
Capping enzyme (CAP). (A) RABV (green) and VSV (gray) CAP domains superimposed. Catalytic residues in the active site (HR and GxxT motifs) and residues participating in Zinc ion coordination are shown for RABV in stick representation. Black dashed circles highlight loops near the active site with different conformations. (B) Zn coordinating residues in RABV CAP (green sticks). Dashed black lines indicate the side-chain atoms involved in metal coordination. (C) RdRp (blue) and CAP (green) domains of RABV (left) and VSV L (right) show similar insertion of the poorly ordered priming loop from CAP into the RdRP active site. The conserved tryptophan from CAP important for transcription initiation from the 3‘ end of the viral genome is the same location for both structures.

**Figure S13:**
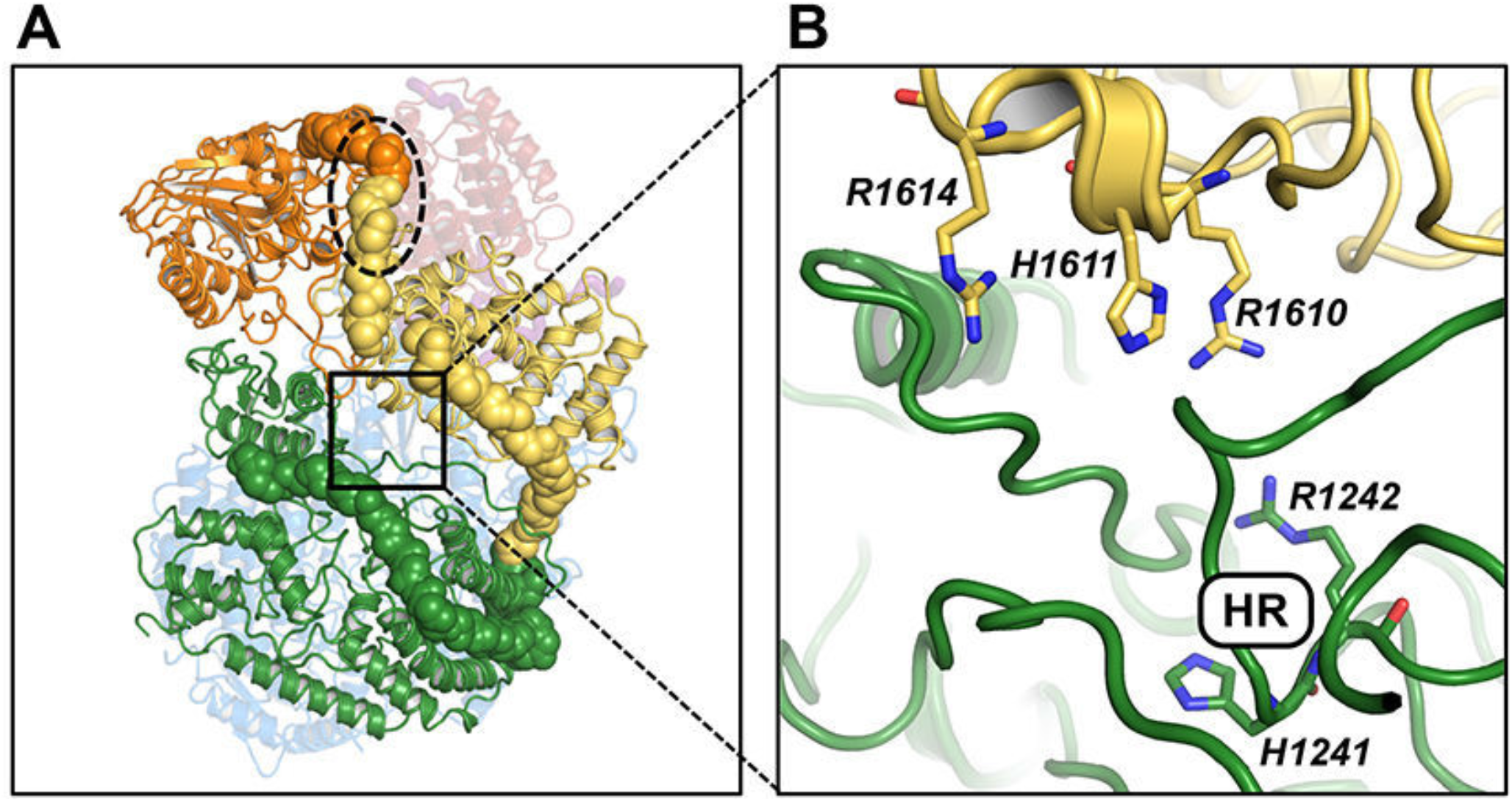
Connector domain (CD). (A) RABV and VSV CDs. (B) Cartoon representation of VSV CAP (green), CD (yellow), and MT (orange) with long linkers between CAP:CD and CD:MT shown as spheres. Black dashed oval, region of the CD:MT linker disordered in the density map. Boxed window and (C), CAP active site at the interface with the CD showing a set of conserved basic residues opposite the HR motif from CAP.

**Figure S14:**
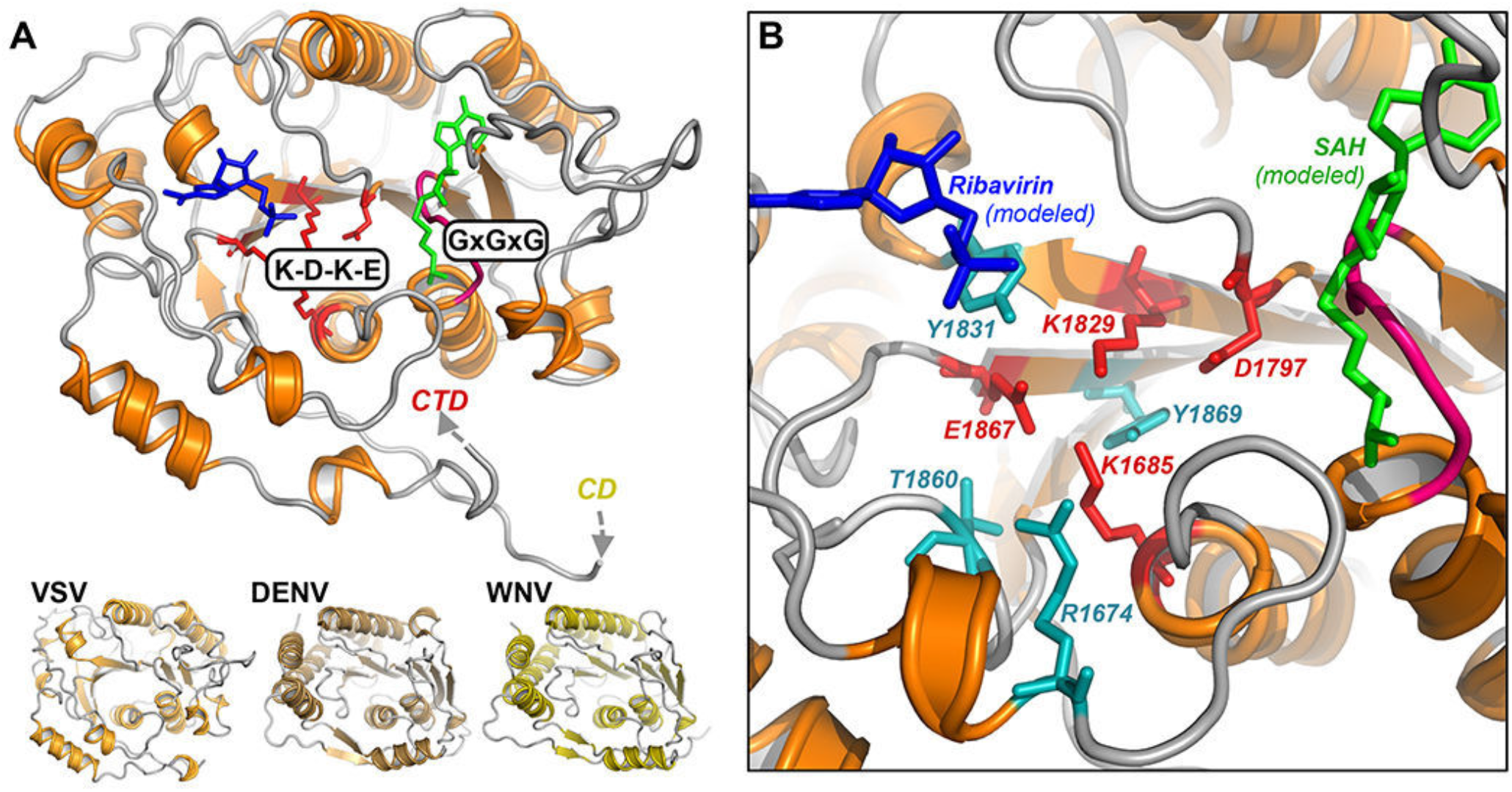
Methyltransferase (MT). (A) RABV (top) and three other dual-function viral MT domains (bottom). Catalytic tetrad residues are in red stick representation; GxGxG motif in pink. The positions of ribavirin (blue) and SAH (green) present in the DENV crystal structure are modeled against the RABV MTase to illustrate likely binding areas for the GTP cap and SAM, respectively. (B) Close-up of the RABV MTase active site, as in (A), showing conserved residues (teal, sticks) that may support catalysis.

